# Ageing conserves and redistributes local geometry in the human structural connectome: an Ollivier-Ricci curvature analysis across the adult lifespan

**DOI:** 10.64898/2026.08.08.743682

**Authors:** Rodrigo Debona, Roger Walz

**Author notes:** Corresponding author: Rodrigo Debona.

## Abstract

Network measures of the ageing connectome are dominated by magnitude: connection strength and density decline, and the topological summaries built on them decline with them. Whether the geometry of the network follows the same course is not known, because the quantities in common use do not separate how strong a connection is from how it sits among the connections around it. We computed the Ollivier-Ricci curvature of every edge in structural connectomes from 307 participants spanning the adult lifespan, a quantity defined by optimal transport between the neighbourhoods of connected regions, and asked how it changes with age. The total geometric separation between within-network and between-network connections did not change across seven decades. Underneath that constancy, individual network pairs moved substantially and in opposite directions, gaining curvature around the salience and ventral attention system and losing it between the control and default mode networks. Curvature and connection strength reached half of their age-related variation almost four decades apart, and a small set of prefrontal nodes moved against the global trend. Ageing appears to conserve the local redundancy of the structural connectome in total while relocating it, on a timescale distinct from that of connection strength.

## INTRODUCTION

The connectome is usually summarised by how strongly its regions are connected. Streamline counts, edge weights, nodal strength, density and the graph statistics built from them all inherit the magnitude of the underlying connections, and in ageing they decline together. Structural connectivity weakens from mid-adulthood, density falls, and global efficiency follows (Betzel et al. 2014; Gong et al. 2009). This convergence is so consistent that it can obscure a question the measures are not designed to answer: whether the arrangement of connections changes on the same schedule as their strength.

The distinction matters because a network can lose weight without changing shape, or change shape while conserving weight, and the two have different consequences for how signals travel. What is missing is a quantity that separates them. Nodal strength is magnitude by definition. Clustering and modularity describe arrangement but are combinatorial constructions, defined by convenience rather than derived from a principle, and they carry no statement about what they control. Efficiency is a summary of path lengths, which are themselves functions of the weights. None of these can say whether the geometry of the network is preserved while its magnitude erodes.

Ollivier-Ricci curvature offers such a quantity. Defined for each edge as a comparison between the cost of optimal transport across it and the length of the edge itself, it asks a local question: do the neighbourhoods of two connected regions overlap, so that alternative short routes exist, or does the edge stand alone as a bottleneck (Ollivier 2009)? Positive curvature marks local redundancy, negative curvature marks a bottleneck, and the correspondence with local clustering is exact enough to have been proved rather than assumed (Jost and Liu 2014). The quantity is not an index invented for networks and later justified; it is the discrete counterpart of a construction in Riemannian geometry, and it inherits results from that setting, including a bound linking curvature to the speed at which diffusion reaches equilibrium.

Curvature has entered neuroimaging along two tracks. Farooq et al. (2019) computed it on structural connectomes to identify robust and fragile regions, and reported, among several demonstrations, that it tracked age. Functional studies have used it to characterise autism (Simhal et al. 2020; Elumalai et al. 2022) and attention-deficit disorder (Chatterjee et al. 2021), and one has addressed ageing directly, contrasting a young group with an older one in functional networks and finding curvature higher in the older group at every graph density examined (Yadav et al. 2023). That study fixes a direction and leaves the course unspecified. A two-group contrast cannot separate a steady decline from one that is flat for decades and then breaks, and it cannot say when geometry moves relative to connection strength; functional connectivity, in addition, does not speak to the anatomical substrate. Following curvature continuously across the adult lifespan in structural connectomes is where the question of whether geometry and magnitude age together can be posed.

The obstacle is partly practical. Exact curvature requires solving one optimal-transport problem per edge, and the analysis is further exposed to a confound that the ageing connectome makes acute. Graphs must be compared at matched size, and proportional thresholding is the standard remedy, but it equalises how many edges are retained without equalising what fraction of each participant’s available edges those represent. When the number of reconstructed edges declines with age, as it does, so does that fraction, and the composition of the retained graph shifts systematically with the variable of interest. Any statistic computed afterwards inherits the shift.

Here we compute exact Ollivier-Ricci curvature for every edge of the structural connectomes of 307 participants spanning the adult lifespan, at fixed graph size, and ask three questions. Whether curvature changes with age, and with what shape. Whether that change follows the same time course as connection strength, or a different one. And whether the change is uniform across the cortex or redistributes geometry among specific systems while leaving its total intact. We address the thresholding confound directly, by equalising the supply of candidate edges before thresholding and correcting for the attenuation this introduces, and we report the effect both before and after. Throughout, we separate what the geometry measures from what it is taken to mean, and we report the analyses that returned nothing alongside those that did not.

## RESULTS

Structural connectomes were reconstructed for 307 Cam-CAN participants (153 female, 154 male; 18.6 to 89.0 years, median 54.3) and parcellated into 200 cortical regions. No participant had an empty parcel. Raw graph density fell with age (*r* = −0.394, *p* = 7.5 × 10^−13^) as did total streamline weight (*r* = −0.565, *p* = 2.9 × 10^−27^), with raw density spanning 0.207 to 0.817 (Annex B, Figure B7). After consistency masking at 60% and proportional thresholding, all 307 graphs retained exactly 2,985 edges, and no participant required edges from outside the group mask (Annex B, Figure B7C).

### Curvature increases with age along a nonlinear course

Mean Ollivier-Ricci curvature rose with age (Spearman *ρ* = 0.329, 95% CI 0.223 to 0.436). Age explained 27.9% of the variance beyond sex, and the course was nonlinear: an F test of the spline block against a linear-only model gave *F* (4, 300) = 13.0, *p* = 9.1 × 10^−10^ (Figure 1A, Table 1). The fitted curve was close to flat until the seventh decade and then rose steeply.

**Figure 1:**
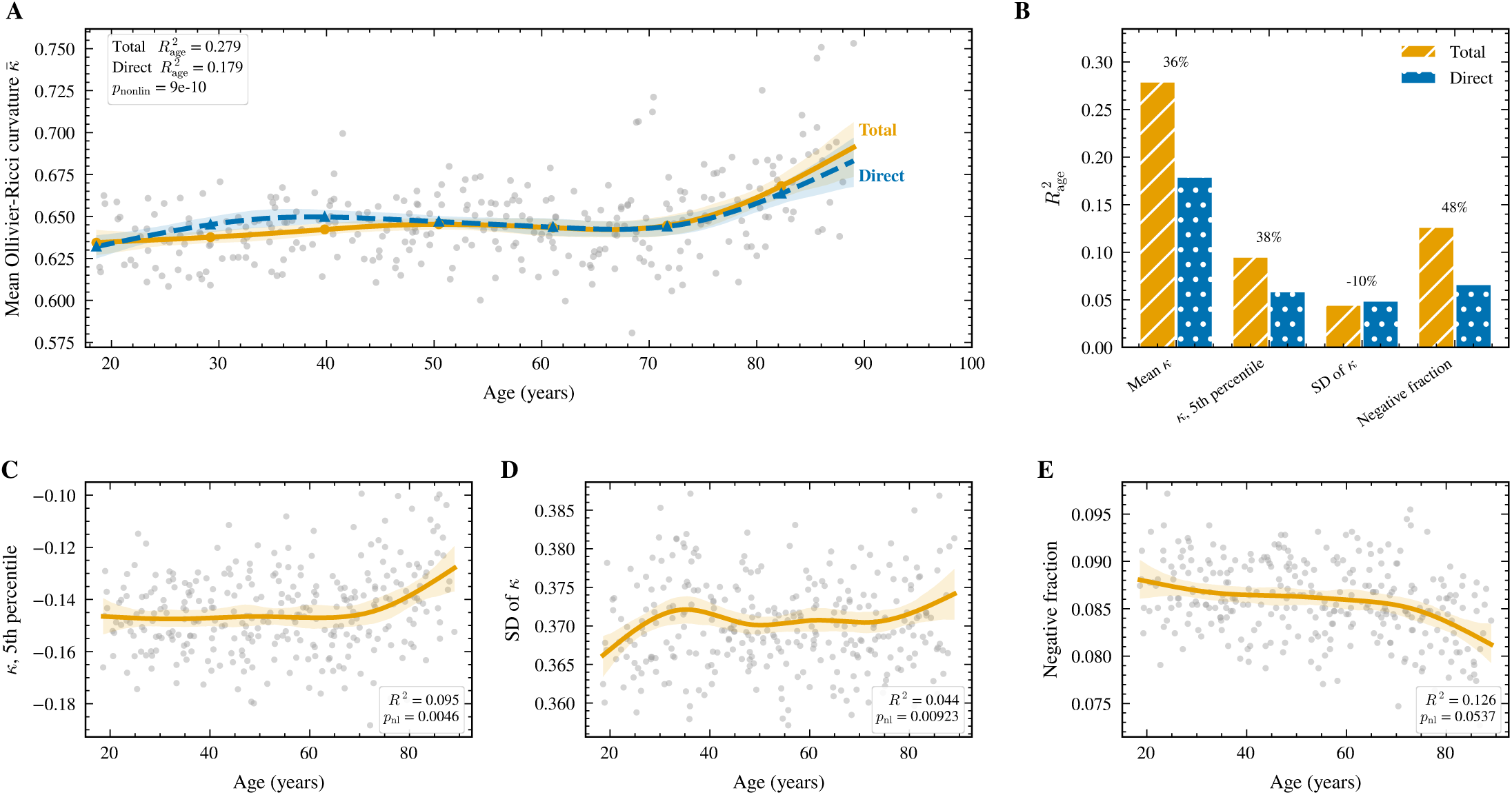
Age-related increase in connectome curvature, and the fraction mediated by magnitude. (A) Mean Ollivier-Ricci curvature against age; orange, total effect adjusted for sex; blue, direct effect adjusted additionally for total streamline weight. Grey points are individual participants and shaded bands are bootstrap 95% confidence intervals. (B) Partial variance in each curvature summary explained by age under both specifications, with the share mediated by magnitude above each pair. (C to E) The remaining curvature summaries under the total-effect specification.

The remaining summaries of the edge curvature distribution moved together with the mean. The fifth percentile rose (*R*^2^ = 0.095, *p*_nonlinear_ = 4.6 × 10^−3^), the dispersion rose (*R*^2^ = 0.044, *p*_nonlinear_ = 9.2 × 10^−3^), and the fraction of negatively curved edges fell (*R*^2^ = 0.126), so the distribution shifted as a whole rather than through a single tail (Figure 1C to 1E).

Adjusting additionally for total streamline weight, which itself declines with age, left 64% of the age variance intact (*R*^2^ = 0.179). Adding total streamline count as a further covariate left *R*^2^ = 0.151 (Figure 1B).

### The increase is independent of thresholding, degree, weighting, fibre length and parcel volume

Five control analyses were run, each removing a different possible source of the association (Table 2, Figure 2).

**Figure 2:**
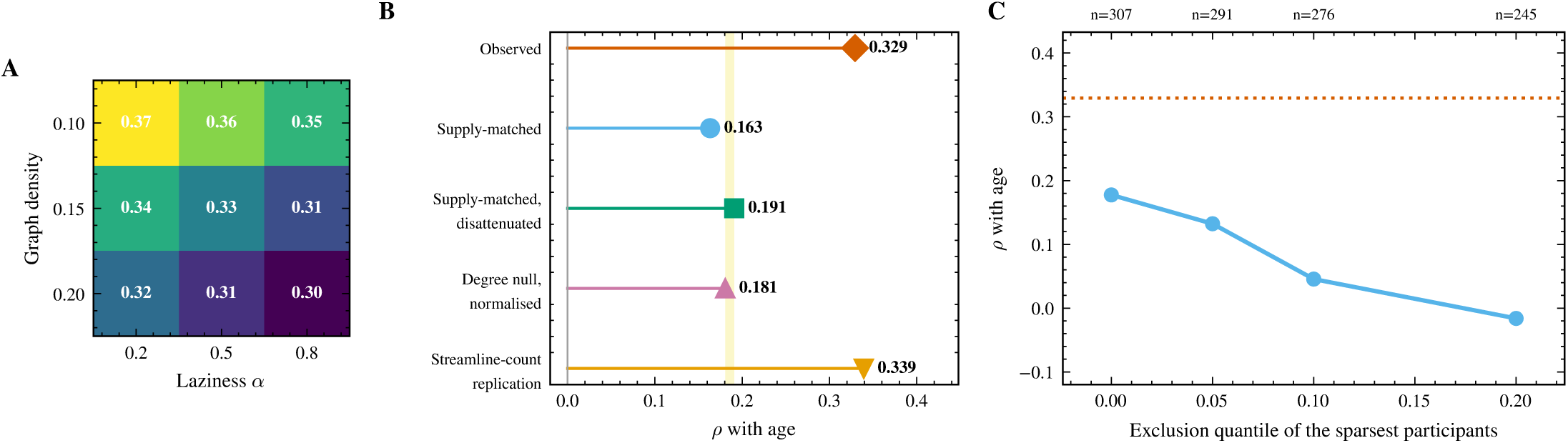
The age effect on curvature survives every control. (A) Spearman correlation between age and mean curvature across a grid of graph density and random-walk laziness; every cell shares the same sign and all *p* < 1.3 × 10^−7^. (B) Effect size under each control; the shaded band marks the interval on which the independent controls converge. (C) Sensitivity of the supply-matched estimate to how aggressively the sparsest participants are excluded.

Equalising the number of candidate edges across participants before thresholding, at a common supply of 4,116 edges, reduced the association to *ρ* = 0.163 across five independent subsamplings. Correcting for the reliability of the subsampled measure (*r*_*xx*_ = 0.732) returned *ρ* = 0.191. Rewiring each connectome while preserving every node degree exactly, and normalising the observed value against five such rewirings per participant, gave *ρ* = 0.181 (*p* = 1.5 × 10^−3^). The two controls are independent and converged on the same estimate.

Across a grid crossing three graph densities with three values of the laziness parameter, all nine cells shared the same sign, with *ρ* between 0.296 and 0.372 and all *p* < 1.3 × 10^−7^ (Figure 2A). Replacing SIFT2 weights with raw streamline counts returned *ρ* = 0.339, and the nodal maps of the age effect from the two weightings correlated at *ρ* = 0.88.

Stratifying edges by group median length, the association held in short edges (*ρ* = 0.354, 11,151 edges), medium edges (*ρ* = 0.371, 4,440 edges) and long edges (*ρ* = 0.297, 4,309 edges), with nonlinearity preserved in all three (all *p* < 5.3 × 10^−8^). The effect was largest in the short and medium strata, the opposite of what a length artefact predicts. Mean length of retained edges fell with age (*r* = −0.310), and adjusting for it left *R*^2^ = 0.111 with *p*_nonlinear_ = 4.9 × 10^−7^ (Annex B, Section B.4).

Mean parcel volume declined with age (*r* = −0.579, *p* = 7.4 ×10^−29^). Adjusting each node for its own parcel volume in native space left the nodal map of age effects essentially unchanged (*ρ* = 0.963 with the unadjusted map; sign preserved at 94.5% of nodes), and 63 of the 67 nodes surviving correction before adjustment also survived after it, including 8 of the 10 decreasing nodes, all 10 of which retained a negative coefficient. Carrying mean parcel volume in the model did not reduce the age effect on mean curvature but increased it, from *R*^2^ = 0.179 to *R*^2^ = 0.220, with nonlinearity preserved (*p* = 1.3 × 10^−9^). The replication with unweighted streamline counts and the most conservative covariate specification are reported in full in Annex B, Sections B.7 and B.8.

### Magnitude and geometry follow different age courses

We standardised both measures within the cohort and modelled their within-participant difference against age. That difference varied with age: *F* (5, 300) = 43.1, *p* < 1 × 10^−16^, *R*^2^ = 0.418 (Figure 3B, Table 3). A permutation test swapping the measure label within participants gave the same conclusion, with no resample of 5,000 exceeding the observed statistic (*p* < 2 × 10^−4^). Residual variance was homogeneous across age (Levene *p* = 0.68).

**Figure 3:**
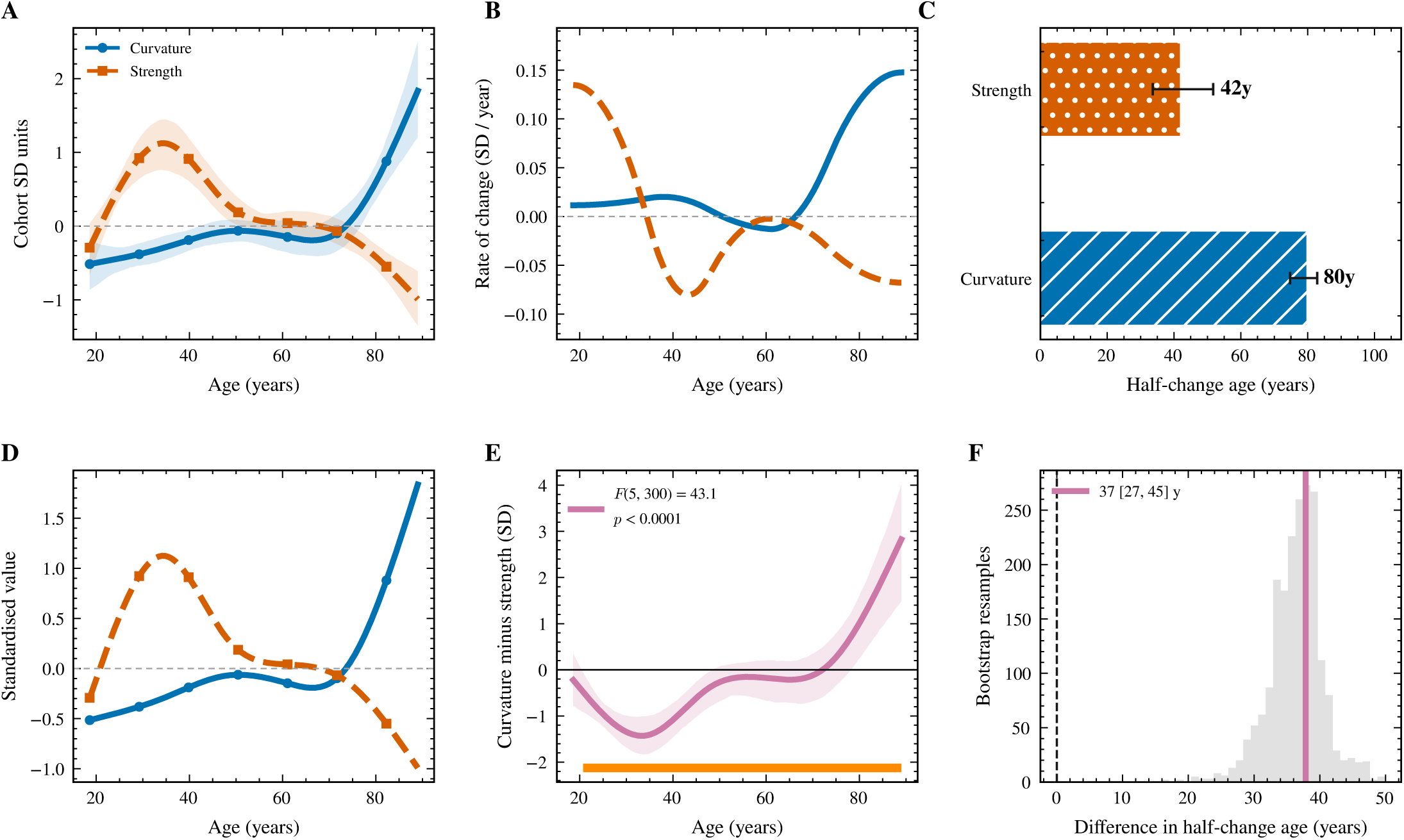
Magnitude and geometry of the structural connectome follow different age courses, and the difference is formally testable. (A) Sex-adjusted spline fits of mean Ollivier-Ricci curvature and of mean nodal strength, each expressed in cohort standard-deviation units; shaded bands are bootstrap 95% confidence intervals. (B) First derivative of each fitted curve, showing when each measure moves fastest. (C) Age at which half of the total variation of each fitted curve has occurred, with bootstrap 95% intervals. (D) The two standardised courses drawn together, which is the comparison the test formalises. (E) The within-participant difference between the two standardised measures. Under the null hypothesis that both follow the same age course up to scale this curve is flat; the bar along the lower axis marks ages at which the confidence band excludes zero. (F) Bootstrap distribution of the difference between the two half-change ages, with the observed value marked and zero dashed. Primary test: *F* (5, 300) = 43.1, p = < 1 × 10^−4^. Half-change at 80 years for curvature and 42 for strength, a difference of 38 years (95% CI 28 to 45). *n* = 307.

Mean curvature reached half of its total variation at 79.8 years (95% CI 72.0 to 80.9) and mean nodal strength at 41.9 years (95% CI 32.8 to 50.8), a separation of 37.9 years (95% CI 27.2 to 44.9). The derivative of each fitted curve places the difference: strength changed fastest during the fourth decade and curvature during the eighth (Figure 3C).

### A prefrontal minority of nodes moves against the global increase

Sixty-seven of 200 nodes showed an age effect surviving false discovery rate correction at 5%, of which 57 increased and 10 decreased (Figure 4A). The nodal map replicated internally: across 200 age-stratified splits of the cohort, maps estimated in the two halves correlated at *ρ* = 0.676 (95% CI 0.598 to 0.747). Nodes of the decreasing set were recovered in a median 78% of 1,000 bootstrap resamples, with 12 selected in at least half of resamples and 5 in at least 80% (Figure 4C). The age course of each node set considered separately, the spatial dispersion of the decreasing set and the bilateral symmetry of the nodal effect map are given in Annex B, Sections B.1 to B.3.

**Figure 4:**
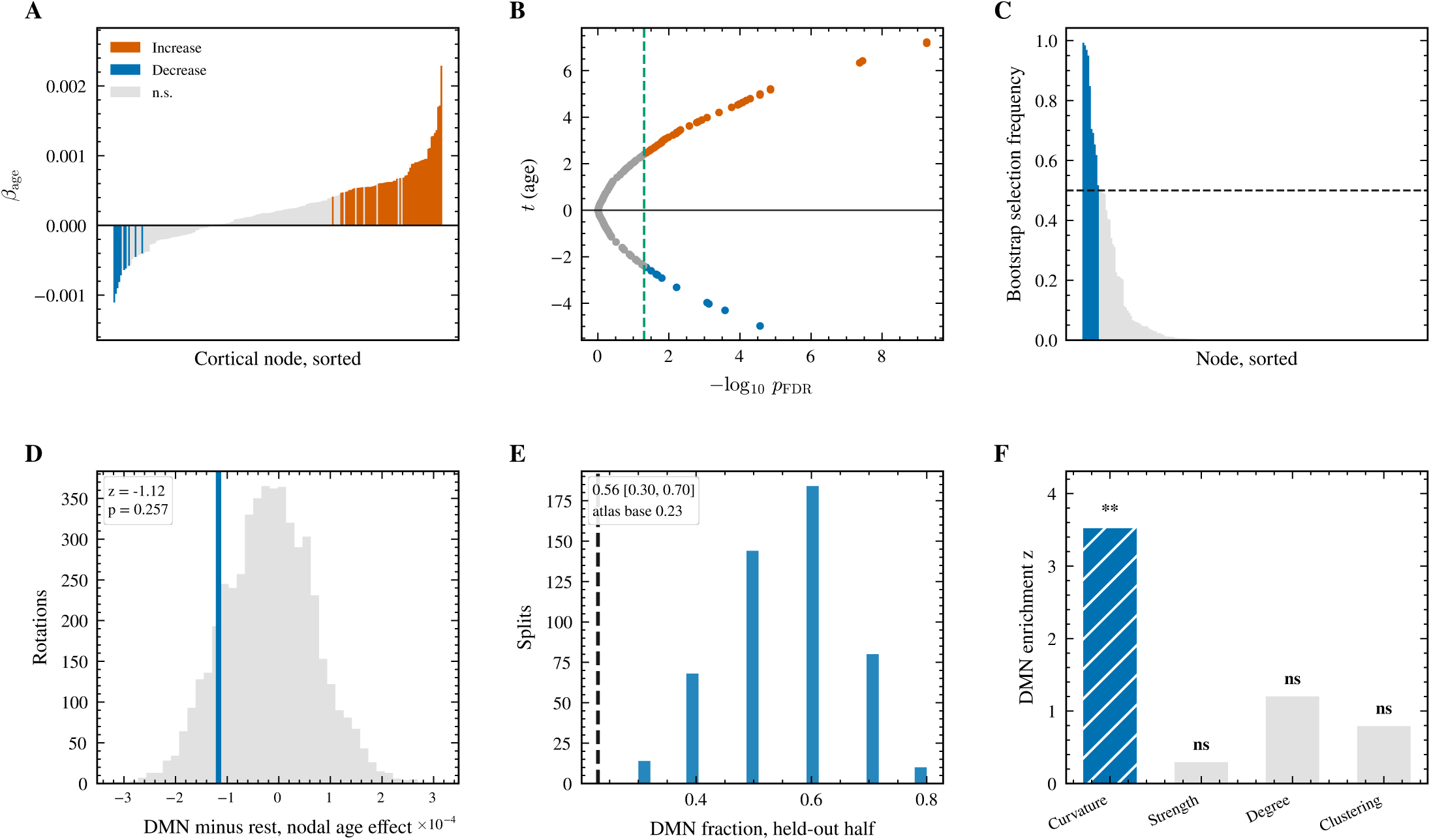
Nodal topography of the age effect, and the controls establishing that its network composition is not an artefact of selection. (A) Age coefficient of nodal curvature for every cortical node, sorted by effect size; colour marks nodes surviving false-discovery-rate correction at 5%. (B) Volcano plot of the same nodal statistics, with the correction threshold dashed. (C) Proportion of 1,000 bootstrap resamples of participants in which each node was retained in the decreasing set. (D) Threshold-free test: the contrast between the mean nodal age effect inside and outside the default mode network, with no node selection and no threshold, against spatially rotated null maps. (E) Out-of-sample test: the decreasing set is defined in one randomly assigned half of the participants and its network composition evaluated in the held-out half, over 500 age-stratified splits; the dashed line is the proportion of default mode nodes in the atlas. (F) Specificity: the identical selection and rotation procedure applied to nodal strength, degree and clustering, with the set size fixed at the value observed for curvature. Abbreviations: DMN, default mode network. Selection and inference use independent participants in panel E, so the enrichment there cannot follow from the selection rule. *n* = 307.

Eight of those 12 nodes belong to the default mode network, two to the control network, one to the salience and ventral attention network and one to the visual network. Ten of the 12 are frontal (Table 5). The dorsal and medial prefrontal subdivision alone contributes five, with two further prefrontal default mode nodes, two prefrontal control nodes and one frontal operculum node. No precuneus, posterior cingulate or parahippocampal node was recovered, and only one lateral temporal node. Nine of the 12 are right hemispheric.

Three analyses established that the composition of this set is specific to curvature and does not depend on how the set was selected (Table 4, Figure 4D to 4F). Applying the identical selection and rotation procedure to nodal strength, degree and clustering, with set size fixed at the value observed for curvature, gave no enrichment of the default mode network for any of them (strength *z* = 0.37, *p* = 0.73; degree *z* = 1.20, *p* = 0.26; clustering *z* = 0.79, *p* = 0.70), against *z* = 3.53, *p* = 1.4 × 10^−3^ for curvature (Annex B, Section B.5). Defining the set in one randomly assigned half of the cohort and evaluating it in the held-out half over 500 age-stratified splits, the pre-selected nodes fell below the rotation null in every one of the 500 splits (*z* = −3.53, SD 0.43), and the default mode fraction of the selected set averaged 0.56 (95% CI 0.40 to 0.80) against an atlas base rate of 0.23. Contrasting the mean nodal age effect inside and outside the default mode network, with no selection at any step, did not reach significance (*z* = −1.12, *p* = 0.257), placing the effect in the tail of the nodal distribution rather than in a shift of the network as a whole.

Spatial nulls throughout were 5,000 rotations of the parcel centroids on the sphere, applied per hemisphere with a common rotation and one-to-one reassignment of parcels.

### Total local redundancy is conserved and redistributed across networks

Collapsing edge curvature into blocks defined by the seven canonical networks gave a symmetric matrix per participant, with within-network edges on the diagonal. Twenty-three of the 28 blocks carried at least 20 retained edges and were modelled; seven survived false discovery rate correction at 5% (Figure 5A, Table 6). The complete set of 23 modelled blocks appears in Annex B, Table B4.

**Figure 5:**
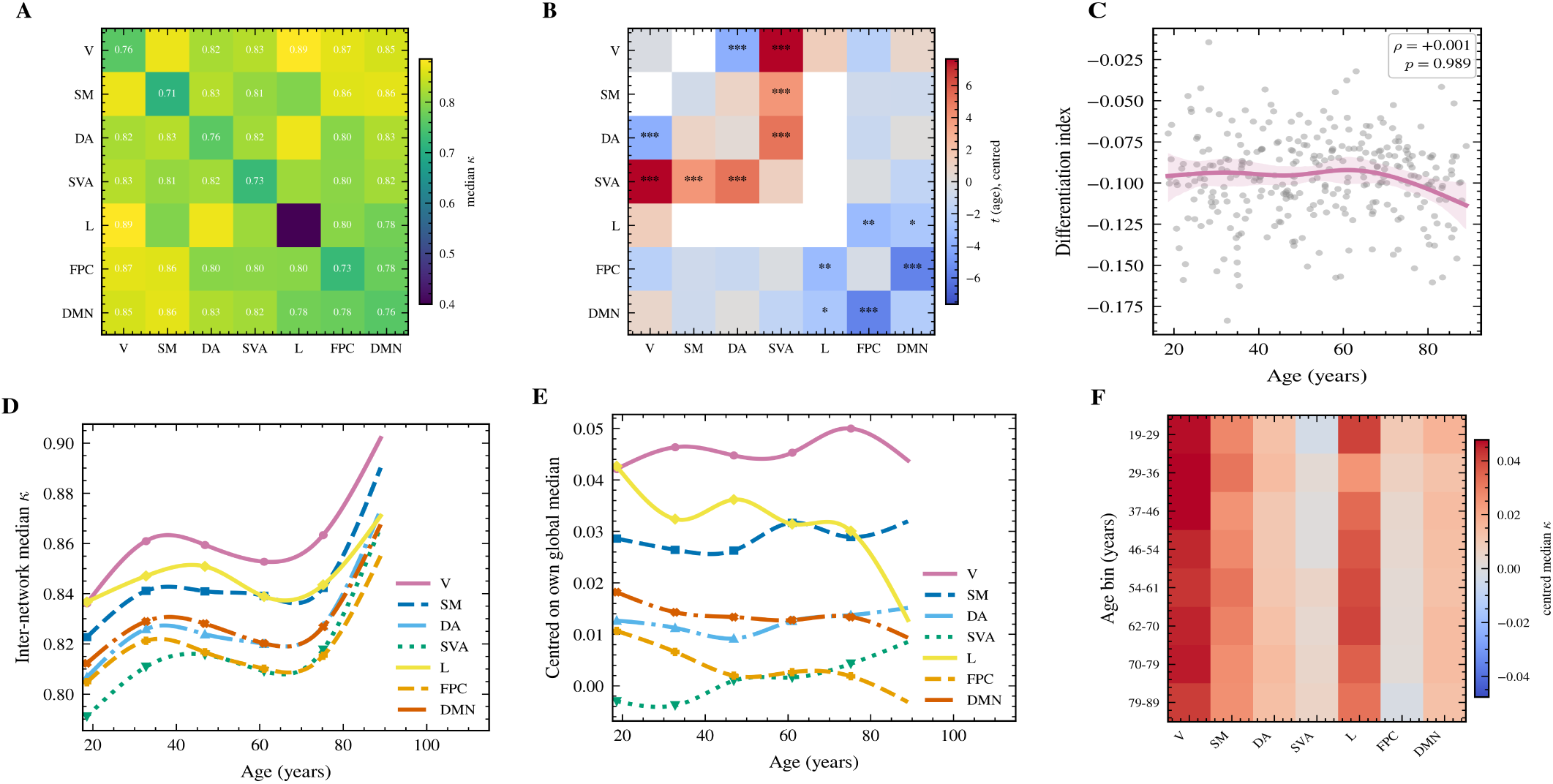
Curvature within and between networks across the adult lifespan. (A) Cohort median of the block curvature matrix. The diagonal holds edges with both endpoints in the same network; off-diagonal cells hold edges joining two different networks. (B) Age effect per block after centring each participant on their own global median curvature, so a cell is coloured by how much faster or slower it changes than the brain as a whole; asterisks mark survival of false-discovery-rate correction. (C) Differentiation index, the median curvature of within-network edges minus that of between-network edges, against age. (D) Median curvature of all edges leaving each network, fitted against age. (E) The same quantity after centring each participant on their own global median, so a curve rising above zero marks a network whose external connections gain curvature faster than the brain as a whole. (F) The centred profile summarised in equal-sized age bins. Abbreviations: V, visual network; SM, somatomotor network; DA, dorsal attention network; SVA, salience and ventral attention network; L, limbic network; FPC, frontoparietal control network; DMN, default mode network. A differentiation index that does not change while individual blocks move in opposite directions is conservation with redistribution, not absence of change. *n* = 307.

The differentiation index, the median curvature of within-network edges minus that of between-network edges, showed no association with age (*ρ* = 0.0008, *p* = 0.99, 95% CI -0.111 to 0.108). Within-network and between-network curvature rose at almost the same rate (*ρ* = 0.225 and *ρ* = 0.254), preserving the geometric separation between the two edge populations. This index was the one quantity sensitive to the summary statistic: computed on block means rather than medians it returned *ρ* = −0.352, although block-level age statistics agreed between the two (*ρ* = 0.73). We therefore report the block matrix as the primary result and the index with both estimates.

Conservation of the global index coexisted with large and opposite changes in individual blocks. Every block involving the salience and ventral attention network gained curvature, most strongly with the visual network (*t* = 7.64, *p*_FDR_ < 1 × 10^−4^), dorsal attention (*t* = 5.09, *p*_FDR_ < 1 × 10^−4^) and somatomotor cortex (*t* = 3.89, *p*_FDR_ = 7.1 × 10^−4^). The largest loss fell between the control and default mode networks (*t* = −5.65, *p*_FDR_ < 1 × 10^−4^), followed by visual to dorsal attention (*t* = −3.77), limbic to control (*t* = −3.20) and limbic to default mode (*t* = −2.50).

The marginal profile, collapsing all edges leaving each network into one value, carried the same polarity (Figure 5C, Table 7). External curvature increased for the salience and ventral attention network (*t* = 5.35, *p*_FDR_ < 1 × 10^−4^) and for somatomotor cortex (*t* = 2.31, *p*_FDR_ = 0.030), and decreased for the control (*t* = −3.40, *p*_FDR_ = 3.0 × 10^−3^), limbic (*t* = −2.83, *p*_FDR_ = 0.012) and default mode (*t* = −2.45, *p*_FDR_ = 0.026) networks. Estimates for the visual and dorsal attention networks did not survive correction (*p*_FDR_ = 0.080 and 0.054). The default mode estimate rests on 951 edges and the control estimate on 718.

### Analyses that returned null results

Curvature was the weakest single predictor of age among the measures tested (cross-validated mean absolute error 10.71 years, against 8.22 for strength and 9.43 for degree). Adding the geometric features to the magnitude features changed the error by 0.21 years, which a sign-flip permutation test on the paired per-participant errors did not distinguish from zero (*p* = 0.20; Annex B, Figure B8).

The nodal age effect correlated with the published principal cortical gradient (*ρ* = −0.216, *p* = 2.2 × 10^−3^ uncorrected), but the correlation did not survive rotation of the effect map (*p* = 0.084 across 5,000 rotations, Figure 6). The ordering of networks along the sensory to transmodal axis is therefore described without inferential support.

**Figure 6:**
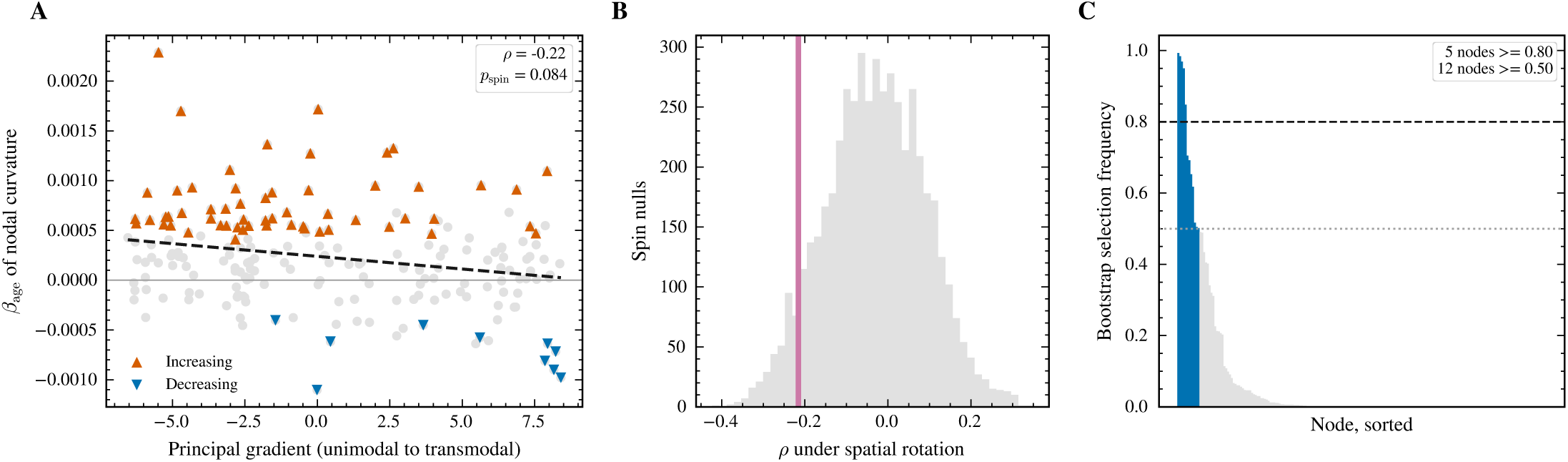
The age effect is not aligned with the principal cortical gradient. (A) Nodal age coefficient against the published principal functional gradient, parcellated into the same regions. (B) Null distribution of that correlation under 5,000 spatial rotations. (C) Bootstrap selection frequency of each node in the decreasing set.

Neither the age effect on mean curvature nor its course interacted with sex (*F* = 0.59, *p* = 0.71). The decreasing and increasing node sets did not differ in degree, participation coefficient, clustering, betweenness or baseline curvature (all *p* > 0.11), although with 10 against 57 nodes these comparisons could detect only effects above *d* = 0.97 at 80% power and are therefore uninformative (Annex B, Figure B9). A test of whether ageing polarised or compressed the existing spread of curvature across nodes also returned nothing; both results are given with their point estimates in Annex B, Section B.10.

## DISCUSSION

Ageing did not erode the local redundancy of the structural connectome so much as relocate it. The total, measured as the geometric separation between within-network and between-network connections, was indistinguishable from constant across seven decades (*ρ* = 0.0008, 95% CI -0.111 to 0.108). Underneath it, individual network pairs changed by as much as eight standard errors and in opposite directions: every block involving the salience and ventral attention network gained curvature, and the largest loss fell between the control and default mode networks. A measure of the average alone would have found nothing, and a measure of magnitude would have found a decline without locating it.

The direction of the global effect is not itself new. Yadav et al. (2023) contrasted young and older adults in functional networks and found curvature higher in the older group at every density they examined; we find the same sign in structural connectomes, in a different cohort and with a continuous design. Agreement across modality, cohort and pipeline is the strongest evidence available that the increase belongs to the ageing network rather than to any one processing chain. What a two-group contrast cannot supply is the course, and the course is where the present result lies: the increase is not steady, and it arrives late.

The claim of conservation sits uneasily beside the dedifferentiation literature. Functional studies from Chan et al. (2014) onward report system segregation declining with age, among the most reproduced findings in the field (Geerligs et al. 2015; Koen and Rugg 2019), and our differentiation index did not. That evidence is almost entirely functional, and its mapping onto structural geometry is not established. Even within it the direction varies: Malagurski et al. (2020) found segregation falling in the default mode, control and salience networks while rising in the limbic. Our index is built from curvature rather than correlation, measuring the separation of two edge populations rather than the coherence of their signals. And the constancy we observe is cancellation rather than absence of change, with visual-to-salience gaining and control-to-default losing at comparable magnitude. Rather than contradicting functional dedifferentiation, the result suggests that the structural substrate reorganises while conserving a quantity those measures do not track.

The minority of nodes moving against the global trend would be weak evidence on its own, since the set was defined by the statistic characterising it. The identical procedure applied to strength, degree and clustering produced no enrichment; defining the set in one half of the cohort and evaluating it in the other placed it below the rotation null in all 500 splits; and the threshold-free contrast, which selects nothing, did not reach significance. We read this as localisation rather than failure: ageing does not displace the default mode network as a whole but drives a subset of its nodes against the prevailing direction, and an average over the network dilutes what lives in its tail. That subset is not spatially clustered (Annex B, Section B.2), so its concentration follows network membership rather than cortical territory.

Those nodes are prefrontal, and the posterior medial core is absent. The set has a nonlinear age course of its own, explaining far less variance than that of the increasing nodes (Annex B, Sections B.1 and B.3); whether it is a subsystem rather than a set unified by sign alone remains to be tested. The anatomy converges with the block analysis, whose largest loss was between the control and default mode networks, both contributions prefrontal: two independent analyses point to the same tissue. This is consistent with prefrontal cortex being affected early and disproportionately in ageing (West 1996), and with the fractionation of the default network into subsystems that need not age together (Andrews-Hanna et al. 2010). Whether it fits a last-in-first-out account is less clear than it once seemed: Robinson et al. (2026) found myelination order a better predictor of white-matter health than order of prenatal emergence, as that account requires, but also that fibre calibre and vascularisation predict susceptibility irrespective of developmental order. We therefore do not read the prefrontal result as evidence for retrogenesis. Nine of the twelve nodes lie in the right hemisphere, an asymmetry we record without interpreting (Cabeza 2002).

The salience and ventral attention network was the consistent recipient: every large gain involved it, and its external curvature rose more than that of any other network. One reading is that as long-range associative connectivity thins, the residual architecture reorganises around this system as a relay, accumulating local redundancy, which aligns with reports that ventral salience connectivity increases with age while its dorsal counterpart declines (Touroutoglou et al. 2018). We offer it as a hypothesis with a test: triangle participation of salience edges should rise with age, by the addition of closing triangles rather than the loss of negatively curved bridges.

Magnitude and geometry reached half of their total age-related variation thirty-eight years apart, and the two measures are not independent, which makes the dissociation harder to obtain rather than easier. Curvature is computed on a graph whose distances derive from the same weights that define strength, so a monotone transformation of strength should produce a similar course, not a different one. It does not. The nonlinearity survives adjustment for total streamline weight at *p* = 5.9 × 10^−10^, and adjustment raises rather than lowers the variance explained by age. Parcel volume behaved the same way, so the unadjusted estimates here are conservative.

### A confound in proportional thresholding

Proportional thresholding equalises how many edges each participant retains, not what fraction of that participant’s available edges those represent, and in this cohort the retained fraction ranged from 0.18 to 0.72 and covaried with age. A participant with many reconstructed edges contributes a selective skeleton of strong connections; a participant with few contributes nearly the whole graph. Same size, different composition, and any variable tracking the number of available edges tracks that difference. Equalising the supply before thresholding, and correcting for the attenuation subsampling introduces, halved the raw association from *ρ* = 0.329 to *ρ* = 0.191. A degree-preserving null, addressing an unrelated concern, converged independently on *ρ* = 0.181, and we take that agreement as the honest estimate.

The safeguard in common use is to sweep the threshold and integrate across it (Elumalai et al. 2022; Yadav et al. 2023), and we swept it as well, over a grid on which every cell carried the same sign. A sweep shows that a result does not depend on one arbitrary density; it cannot show that graphs compared at that density were cut to the same depth into their own weight distributions, because depth is set by supply, which the sweep leaves untouched. The two controls are orthogonal, and only the second moved our estimate. The exposure is not specific to curvature: any statistic computed after proportional thresholding inherits it whenever the number of reconstructed edges covaries with the variable of interest, as in ageing (Betzel et al. 2014). Heuvel et al. (2017) described a related bias in functional networks in terms of overall connectivity strength; the formulation in terms of the fraction of available edges, and its consequences for structural networks, has not to our knowledge been stated. Matching a whole distribution rather than a summary of it is the move already made in harmonisation (Zhou, Fischl, and Aganj 2025). Consistency-based thresholding avoids the problem by a different route (Roberts et al. 2017); supply matching offers a check applicable to existing pipelines unchanged.

### What the geometry does not do

Curvature was the weakest single predictor of age among the measures tested, and adding it to magnitude did not significantly improve prediction. The contribution here is mechanistic rather than predictive. The nodal effect map correlated with the principal cortical gradient, but the correlation did not survive spatial rotation, so we describe the ordering of networks along the sensory-to-transmodal axis without claiming alignment. Spatial autocorrelation inflates such correlations substantially (Markello and Mišić 2021), and we would rather forgo the claim than make it on an uncorrected test.

### Limitations

The design is cross-sectional: the trajectories are age-related differences rather than within-person change, cohort effects cannot be separated from ageing, and the late inflection requires longitudinal confirmation. Tractography recovers connections with substantial false-positive and false-negative rates and underestimates long connections (Maier-Hein et al. 2017; Schilling et al. 2019); we recovered the effect within each fibre-length stratum (Annex B, Section B.4), but the limitation stands, and curvature depends on the local distribution of edge weights rather than their sum, inheriting that uncertainty at a finer grain than nodal measures do. Microscopy-informed estimation of fibre orientations is beginning to constrain reconstruction where it is weakest (Zhu et al. 2026), and is the natural substrate on which to re-test this. Head motion was not available in this release, though streamline count, which partially proxies it, was a covariate. The analysis uses one parcellation at one resolution, and the comparison of decreasing and increasing node sets on nodal properties detected only effects above *d* = 0.97, so its null is not evidence of equivalence. Finally, the curvature-robustness correspondence rests on an analogy with the fluctuation theorem rather than on a theorem about graphs (Farooq et al. 2019), and we have kept our claims to what the geometry measures directly.

### Conclusion

Local redundancy in the ageing structural connectome is conserved in total and redistributed in detail, on a timescale distinct from connection strength. The geometry moves late, moves against the global trend in a set of prefrontal nodes, and accumulates around the salience system. None of this is visible to measures of magnitude, and the effect size is modest by design: half of the raw association was an artefact of thresholding, and we report what survived.

## MATERIALS AND METHODS

### Participants, acquisition and connectome reconstruction

Diffusion-weighted images were taken from the Cambridge Centre for Ageing and Neuroscience (CamCAN) Stage∼2 CC700 release (Shafto et al. 2014; Taylor et al. 2017). Data were acquired on a 3∼T Siemens TIM Trio using a twice-refocused spin-echo EPI sequence (Reese et al. 2003) sampling 30 gradient directions at each of *b* = 1000 and *b* = 2000∼s,mm^−2^ plus three non-diffusion-weighted volumes at 2∼mm isotropic resolution; full sequence parameters are listed in Annex∼C, Table∼C1. The present analysis used 307 participants (age range 18.6–89.0 years, median 54.3, mean 53.7, SD 19.5; 153 women and 154 men), selected to be balanced across the adult lifespan. Preprocessing was carried out in MRtrix3 3.0.8, which called FSL 6.0.7.22 and ANTs 2.6.5 where indicated. Each series was denoised by Marchenko–Pastur PCA before any interpolating operation (Veraart et al. 2016), corrected for Gibbs ringing by the local subvoxel-shift method with 20 shifts (Kellner et al. 2016), and corrected for eddy-current distortion and head motion with FSL `eddy` using a linear second-level model, outlier detection and replacement, and post-eddy shell alignment disabled. The protocol contains no reverse phase-encoded volumes, so susceptibility distortion was estimated from a synthetic undistorted *b* = 0 image predicted from the participant’s T1w scan by Synb0-DisCo (Schilling et al. 2019, 2020). The synthetic image and the measured mean *b* = 0 image were passed to TOPUP as a pair with, respectively, zero and measured total readout time, and the resulting field was applied to the eddy-corrected series with Jacobian intensity modulation. Residual B1 inhomogeneity was removed with N4 (Tustison et al. 2010), and the brain mask was obtained by rigidly registering an HD-BET T1w mask (Isensee et al. 2019) to the mean *b* = 0 image.

Because the acquisition samples two non-zero shells, tissue response functions for white matter, grey matter and CSF were estimated with the unsupervised Dhollander algorithm (Dhollander, Raffelt, and Connelly 2016; Dhollander et al. 2019), and fibre orientation distributions were reconstructed by multi-shell multi-tissue constrained spherical deconvolution (Jeurissen et al. 2014), followed by multi-tissue log-domain intensity normalisation (Raffelt et al. 2017). Anatomically constrained tractography (Smith et al. 2012) used a five-tissue-type image generated from the T1w scan with FSL FAST and FIRST, rigidly coregistered to the diffusion data through a 6-degree-of-freedom, mutual-information registration of the skull-stripped T1w to the mean *b* = 0 image. Five million streamlines were generated per participant with the second-order integration over fibre orientation distributions algorithm (iFOD2; Tournier et al., 2010), seeded from the grey-matter/white-matter interface, with an FOD amplitude cutoff of 0.06, a step size of 0.625∼mm, a maximum angle of 45^∘^ per step, length limits of 5 and 250∼mm, backtracking enabled and streamline endpoints cropped at the interface. Streamline weights that make the reconstruction quantitatively consistent with the FOD lobe integrals were then estimated with SIFT2 (Smith et al. 2015); no streamlines were discarded.

Network nodes were the 200 cortical parcels of the Schaefer 7-network parcellation (Schaefer et al. 2018), mapped onto the Yeo 7-network solution (Yeo et al. 2011). The parcellation was obtained in each participant’s own anatomical space by projecting the fsaverage annotation onto the individual surface reconstruction and filling the cortical ribbon with FreeSurfer 8.2.0 (Fischl 2012), which removes any dependence on a template-to-subject nonlinear warp. FreeSurfer cortical codes were remapped to connectome indices 1–100 for the left hemisphere and 101–200 for the right, and the label volume was carried into diffusion space in world coordinates by applying the T1w-to-diffusion rigid transform once, with nearest-neighbour interpolation onto the mean *b* = 0 grid. Registration source images were verified to be skull-stripped before use, and each parcellation was checked for completeness against the expected 200 parcels. Label identity was verified separately in every participant against the fsaverage colortable, the observed FreeSurfer code ranges and the laterality of each parcel centroid; no participant failed any of these checks.

Streamline endpoints were assigned to parcels by a radial search of up to 4∼mm from each endpoint; a streamline was retained only when both endpoints reached a parcel. The structural connectivity matrix was defined as the sum of SIFT2 weights over the streamlines connecting each pair of parcels, symmetric and with zero diagonal, yielding one 200×200 matrix per participant. Matrices weighted by streamline count, mean streamline length, mean inverse length and the along-streamline mean of each available scalar map were produced from the same endpoint assignments and retained for sensitivity analyses. Per participant we recorded the SIFT2 proportionality coefficient (2.47×10^−4^ ±4.78×10^−5^), the fraction of streamlines contributing to an inter-parcel edge (0.501 ± 0.042) and the number of parcels absent from the resampled parcellation, which was zero in every retained participant. Five of the 312 datasets were excluded before analysis: three whose assignment fraction lay more than three standard deviations from the cohort mean, and two for which `tck2connectome` failed.

The full processing chain, naming every command-line option and the estimator behind each step, is given in Annex C.

### Edge construction and harmonisation

Two properties of the SIFT2-weighted matrices required harmonisation before any comparison across participants. Graph density and total streamline weight both declined with age, and raw density ranged from 0.207 to 0.817 across the sample. Three operations were applied in the following order.

First, a group consistency mask retained node pairs connected in at least 60% of participants. Second, the maximum spanning tree of each masked matrix was forced into the retained set, guaranteeing a connected graph for every participant. Third, the strongest remaining edges inside the mask were added until an identical edge count was reached, corresponding to a density of 0.15, that is 2,985 of the 19,900 possible pairs. All 307 graphs reached exactly this count, and no participant required edges from outside the consistency mask.

Ollivier-Ricci curvature requires a cost on each edge rather than an affinity. We converted the SIFT2 weight *w*_*ij*_ to a distance by *d*_*ij*_ = 1/*w*_*ij*_. The logarithmic alternative *d*_*ij*_ = − log(*w*_*ij*_/*w*_max_) was rejected because it sends the strongest edge to zero distance, at which the curvature ratio defined below diverges; in this dataset it produced mean curvatures on the order of −10^5^.

### Ollivier-Ricci curvature

For each node *x* we defined the one-step lazy random-walk measure

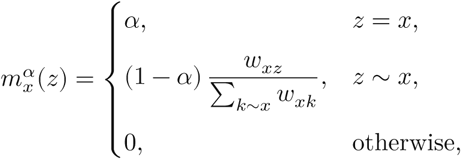

where *α* ∈ [0, 1) is the laziness, the probability that the walker remains in place, and mass is distributed over neighbours in proportion to edge affinity. The Ollivier-Ricci curvature of an edge (*x*, *y*) is

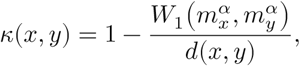

where *W*_1_ denotes the Wasserstein-1 distance, that is the minimum cost of transporting one measure into the other, with cost given by the shortest-path distance computed on the edge distances *d*_*ij*_(Ollivier 2009). The quantity compares the cost actually paid by optimal transport against the cost of moving the mass along the edge itself. Positive curvature indicates that the two neighbourhoods overlap, so that alternative short routes exist; negative curvature indicates that the edge is a bottleneck through which transport is forced. By construction *n* ≤ 1.

Each edge therefore requires one optimal-transport problem. These were solved exactly by linear programming with the Python Optimal Transport library (`ot.emd`; Flamary et al. (2021)) rather than by entropic approximation, which biases the transport cost upward and does so non-uniformly with neighbourhood size. The laziness was set to *α* = 0.5 and varied in a sensitivity analysis (Results).

The implementation was validated against the three graph families for which *κ* has a closed form, the complete graph, the long cycle and the infinite regular tree, with agreement exact to 2.2 × 10^−16^ over twelve parameter combinations. Derivations and the full validation are given in Annex A.

Nodal curvature was the mean over edges incident to a node. Distribution-level summaries were the mean, the fifth percentile, the standard deviation and the fraction of negatively curved edges, computed over the edges of each participant’s graph.

### Network-block curvature

Edges were assigned to blocks defined by the seven canonical networks, giving a symmetric 7 × 7 matrix per participant with within-network edges on the diagonal. Each block was collapsed by its median rather than its mean: the edge curvature distribution is left-skewed and block sizes differ by a factor of four, so a block mean is dominated by a few bridges. The mean was retained as a sensitivity analysis.

Because global curvature increases with age, blocks were additionally centred within participant on that participant’s global median curvature, which isolates blocks changing faster or slower than the brain as a whole. Blocks carrying fewer than 20 retained edges were excluded from modelling rather than being modelled and then corrected, leaving 23 of the 28 blocks. A marginal profile was obtained by collapsing all edges leaving a given network into a single value per participant.

The differentiation index was defined as the median curvature of within-network edges minus that of between-network edges, a scalar measuring the geometric separation of the two edge populations.

### Statistical modelling

Age trajectories were fitted with natural cubic splines on five degrees of freedom, with knots at the sample sextiles, using ordinary least squares. Confidence bands were obtained by bootstrapping participants, with 800 resamples for display and 5,000 for reported intervals. Nonlinearity was tested by an *F* test of the spline block against a model retaining the linear age term, so that the numerator carries four degrees of freedom.

Total streamline weight declines with age and lies on the causal path between age and connectome geometry. It is therefore a mediator rather than a confounder, and adjusting for it estimates a direct rather than a cleaner total effect. We report both specifications throughout: the total effect adjusts for sex alone, and the direct effect adjusts additionally for total streamline weight. Variance attributable to age is reported as the partial *R*^2^ of the spline block over and above the covariates, not the *R*^2^ of the full model.

The dissociation between the age courses of curvature and of magnitude was tested on the within-participant difference of the two cohort-standardised measures. Under the null hypothesis that both follow the same age course up to scale, that difference does not depend on age; the test is therefore an *F* test of the spline block on the difference, against a model with no age term, and the numerator carries five degrees of freedom. A distribution-free counterpart flipped the sign of each participant’s difference over 5,000 resamples. Effect size was expressed as the age at which half of the total variation of each fitted curve has occurred, with the difference between the two and its bootstrap interval reported. Contrasting the coefficients of a fitted polynomial is an alternative route to the same comparison (Seraji et al. 2026); the half-change age was preferred because it is expressed in years and is therefore interpretable and comparable across measures on different scales.

Nodal effects were estimated by ordinary least squares with age, sex and total streamline weight as predictors, and corrected across the 200 nodes by the Benjamini-Hochberg procedure at *q* = 0.05 (Benjamini and Hochberg 1995). Reproducibility of the nodal map was assessed by 200 age-stratified splits of the cohort, correlating the maps estimated in the two halves, and stability of the decreasing node set by 1,000 bootstrap resamples of participants, recording the proportion of resamples in which each node was retained.

Cross-validated prediction of age used ridge regression with the penalty chosen by internal generalised cross-validation, in ten outer folds. Incremental validity was tested on the paired per-participant absolute errors of two nested feature sets by flipping the sign of each participant’s difference over 20,000 resamples, which respects the pairing that pooling across participants would discard.

### Null models

#### Threshold-depth control

Proportional thresholding equalises how many edges each participant retains but not the fraction of that participant’s positive edges those represent, and that fraction covaried with age, ranging from 0.18 to 0.72. Before thresholding, the positive edges of every participant were therefore subsampled uniformly, ignoring weight, to a common supply of 4,116 candidates, set by the sparsest participant. Sampling proportional to weight would have reintroduced the selectivity the control removes. Five independent subsamplings were run. Because subsampling discards information and therefore attenuates any correlation on its own, the resulting estimate was correc_√_ted for attenuation by the reliability between independent repetitions, *r*_*xx*_ = 0.732, as 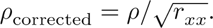

#### Degree-preserving null

Each connectome was rewired by double edge swaps that preserve every node degree exactly and carry edge weights with the edges, with ten swaps attempted per edge and swaps rejected if they would disconnect the graph. Five rewirings were generated per participant, and the observed mean curvature was normalised as 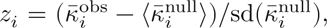 which asks how much curvature a brain carries beyond what its degree sequence alone would produce.

#### Spatial null

Spatial autocorrelation was accounted for by rotating parcel centroids on the sphere (Alexander-Bloch et al., 2018; Váša et al., 2018). Centroids were taken from the Schaefer release in MNI space and projected onto the unit sphere. Rotations were generated with `brainnulls`, using the one-to-one reassignment of Váša et al. (2018), so that each null map is a strict permutation of the observed one and the marginal distribution is preserved exactly. The two hemispheres were rotated with the same sequence of rotations, applied independently within each hemisphere. Five thousand rotations were used throughout. A diagnostic confirming that the nulls retain spatial autocorrelation is reported in Annex A.

#### Circularity controls

The set of nodes whose curvature decreases with age was defined by the curvature statistic itself, so its network composition cannot be tested on the same data without circularity. Three analyses addressed this, each breaking it by a different route. A threshold-free contrast compared the mean nodal age effect inside and outside a network with no selection, no threshold and no correction step. An out-of-sample test defined the set in one randomly assigned half of the cohort and evaluated it in the held-out half over 500 age-stratified splits. A specificity test applied the identical selection and rotation procedure to nodal strength, degree and clustering, with set size fixed at the value observed for curvature.

### Control analyses

#### Parameter sensitivity

The two free parameters of the estimator, graph density and laziness, were crossed in a three-by-three grid spanning densities of 0.10, 0.15 and 0.20 and laziness of 0.2, 0.5 and 0.8.

#### Weighting

The whole pipeline was repeated with raw streamline counts in place of SIFT2 coefficients, using the same consistency mask and density.

#### Fibre length

Tractography underestimates long connections and ageing reduces them, so edges were stratified into terciles by the group median length of each edge, taken from the streamline length matrices, and the analysis was repeated within each stratum. Mean length of the retained edges was also carried as a covariate.

#### Parcel volume

Cortical parcels shrink with age and smaller parcels capture fewer streamline endpoints, a confound that no graph-level control removes because all of them act after the matrix is built. Parcel volumes were computed by counting voxels of each cortical label in the participant’s own parcellation volume, in native conformed space at 1 mm isotropic resolution, so that volume is expressed in the same space where endpoint assignment occurred. Each node was then modelled with its own parcel volume as an additional covariate. Volumes were left unnormalised by intracranial volume, since the confound to be controlled operates in native space.

#### Cortical gradient

The nodal age-effect map was compared against the principal functional gradient (Margulies et al. 2016) as distributed with `brainspace` (Vos de Wael et al. 2020), parcellated into the same 200 regions on the shared conte69 surface. The published gradient was used rather than one derived from this cohort, which would have introduced circularity. Label order was verified by confirming that the parcellated gradient orders visual and somatomotor cortex at the unimodal extreme and the default mode network at the transmodal extreme.

### Software and reproducibility

Analyses were run in Python 3.11 with NumPy 2.4, SciPy 1.17, NetworkX 3.6, scikit-learn 1.8 and POT 0.9.7. Curvature, harmonisation, null models and statistical helpers are provided as a single library, and every analysis is a script that reads only cached intermediates, so that each figure and table can be regenerated without recomputing curvature. Exact curvature took approximately 1.1 s per participant at a density of 0.15 with 200 nodes. The analysis code is available at Zenodo (Debona and Walz 2026), together with 400 synthetic connectivity matrices generated by a network trained on the study matrices. The synthetic matrices carry the marginal and topological properties the pipeline depends on, so every step can be executed end to end from the repository alone; they are not the study data and no reported result should be reproduced from them.

**Figure 7:**
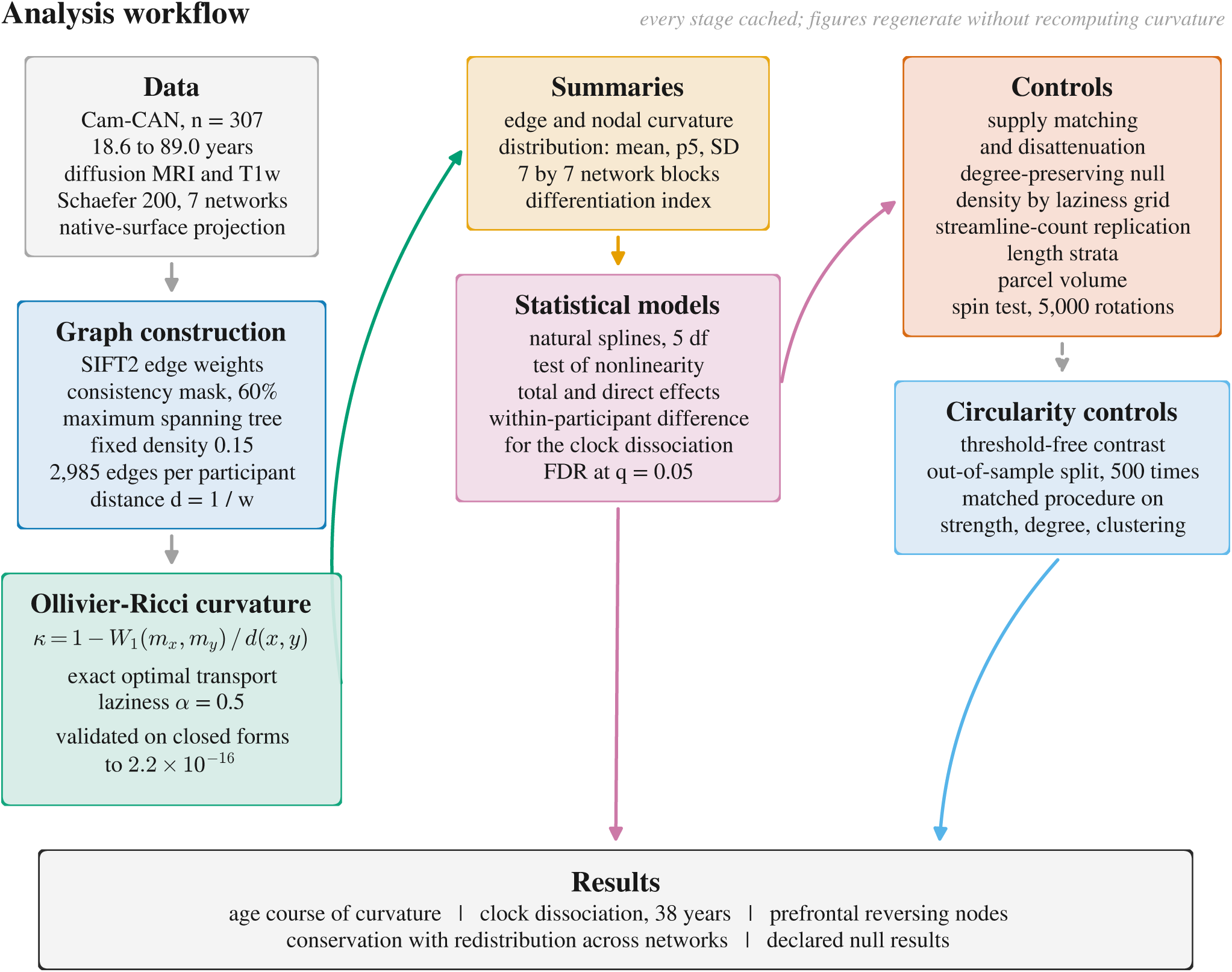
Analysis workflow. Every stage writes cached intermediates, so that figures and tables regenerate without recomputing curvature. Boxes on the left are the construction of the object of study; the centre column is estimation and modelling; the right column is the control analyses, each targeting a distinct way in which the association could arise without geometry changing.

The Cam-CAN data themselves cannot be redistributed by us, since the data access agreement we accepted reserves distribution to the Cam-CAN team. They are available free of charge from https://cam-can.mrc-cbu.cam.ac.uk/dataset/ following the access procedure described there, and Annex C documents every step from the released diffusion data to the matrices analysed here, so that they can be reconstructed exactly.

## Authors, affiliations and declarations

### Authors

**Rodrigo Debona**^1^*, **Roger Walz**^2^

^1^ Instituto Nacional de Neurociência Translacional, Universidade Federal do Rio de Janeiro, Rio de Janeiro, RJ, Brazil.

^2^ Departamento de Clínica Médica, Universidade Federal de Santa Catarina, Florianópolis, SC, Brazil.

* Corresponding author: Rodrigo Debona

## Acknowledgements

We thank the volunteers who took part in the Cam-CAN study, and the Cam-CAN team for making data of this quality openly available to the research community.

Data collection and sharing for this project was provided by the Cambridge Centre for Ageing and Neuroscience (Cam-CAN). Cam-CAN funding was provided by the UK Biotechnology and Biological Sciences Research Council (grant number BB/H008217/1), together with support from the UK Medical Research Council and University of Cambridge, UK.

R.D. thanks CAPES for its support of graduate training in Brazil. R.W. thanks CAPES and CNPq for their support.

## Funding

This work received no specific grant from any funding agency in the public, commercial or not-for-profit sectors.

## Competing interests

The authors declare no competing interests. The work was carried out independently, using data made openly available by the Cam-CAN team following a formal data access request and in accordance with the terms of that agreement.

## Data and code availability

The analysis code is deposited at Zenodo [Debona and Walz (2026); https://doi.org/10.5281/zenodo.21771265], together with 400 synthetic connectivity matrices generated by a network trained on the study matrices. The synthetic matrices carry the marginal and topological properties the pipeline depends on, so every step can be run end to end from the repository alone; they are not the study data and are not a substitute for it.

The Cam-CAN data cannot be redistributed by us. The data access agreement we accepted, and which we continue to honour, reserves distribution to the Cam-CAN team. The data are available free of charge from https://cam-can.mrc-cbu.cam.ac.uk/dataset/ following the access procedure described there. Annex C of the Supplementary Materials documents every step from the released diffusion data to the weighted connectivity matrices analysed here, including software versions and command-line options, so that a reader with access to the data can reconstruct the matrices exactly.

## Author contributions

Following the CRediT taxonomy.

**R.D.**: conceptualization, methodology, software, validation, formal analysis, data curation, visualization, writing (original draft), writing (review and editing).

**R.W.**: conceptualization, methodology, formal analysis, supervision, writing (review and editing).

## Use of artificial intelligence

Artificial intelligence (Claude, Opus 5, Anthropic Inc., <www.claude.ai>) was used at several stages of this work. It contributed to the writing of the analysis code, assisted with the literature review, and was used for linguistic revision of the manuscript, the latter being warranted because the authors are not native speakers of English. It was also used in formatting the manuscript and compiling the LaTeX source. All scientific decisions, including the framing of the research question, the selection and interpretation of analyses, and the final content of the manuscript, were made by the authors, who take full responsibility for the work. All references cited were read and verified by the authors.

## Notes

https://zenodo.org/records/21771265

### Competing Interest Statement

The authors have declared no competing interest.

## References

Andrews-Hanna, Jessica R., Jay S. Reidler, Jorge Sepulcre, Renee Poulin, and Randy L. Buckner. 2010. “Functional-Anatomic Fractionation of the Brain’s Default Network.” Neuron 65 (4): 550–62. 10.1016/j.neuron.2010.02.005.

Benjamini, Yoav, and Yosef Hochberg. 1995. “Controlling the False Discovery Rate: A Practical and Powerful Approach to Multiple Testing.” Journal of the Royal Statistical Society: Series B 57 (1): 289–300.

Betzel, Richard F., Lisa Byrge, Ye He, Joaquín Goñi, Xi-Nian Zuo, and Olaf Sporns. 2014. “Changes in Structural and Functional Connectivity Among Resting-State Networks Across the Human Lifespan.” NeuroImage 102: 345–57. 10.1016/j.neuroimage.2014.07.067.

Cabeza, Roberto. 2002. “Hemispheric Asymmetry Reduction in Older Adults: The HAROLD Model.” Psychology and Aging 17 (1): 85–100. 10.1037/0882-7974.17.1.85.

Chan, Micaela Y., Denise C. Park, Neil K. Savalia, Steven E. Petersen, and Gagan S. Wig. 2014. “Decreased Segregation of Brain Systems Across the Healthy Adult Lifespan.” Proceedings of the National Academy of Sciences 111 (46): E4997–5006. 10.1073/pnas.1415122111.

Chatterjee, Tanima, Réka Albert, Stuti Thapliyal, Nazanin Azarhooshang, and Bhaskar DasGupta. 2021. “Detecting Network Anomalies Using Forman–Ricci Curvature and a Case Study for Human Brain Networks.” Scientific Reports 11: 8121. 10.1038/s41598-021-87587-z.

Debona, Rodrigo, and Roger Walz. 2026. “Analysis Code and Synthetic Structural Connectomes for “Ageing Conserves and Redistributes Local Geometry in the Human Structural Connectome”.” Zenodo. 10.5281/zenodo.21771265.

Dhollander, Thijs, Remika Mito, David Raffelt, and Alan Connelly. 2019. “Improved White Matter Response Function Estimation for 3-Tissue Constrained Spherical Deconvolution.” In Proceedings of the International Society for Magnetic Resonance in Medicine, 27:555.

Dhollander, Thijs, David Raffelt, and Alan Connelly. 2016. “Unsupervised 3-Tissue Response Function Estimation from Single-Shell or Multi-Shell Diffusion MR Data Without a Co-Registered T1 Image.” In ISMRM Workshop on Breaking the Barriers of Diffusion MRI, 5.

Elumalai, Pavithra, Yasharth Yadav, Nitin Williams, Emil Saucan, Jürgen Jost, and Areejit Samal. 2022. “Graph Ricci Curvatures Reveal Atypical Functional Connectivity in Autism Spectrum Disorder.” Scientific Reports 12: 8295. 10.1038/s41598-022-12171-y.

Farooq, Hamza, Yongxin Chen, Tryphon T. Georgiou, Allen Tannenbaum, and Christophe Lenglet. 2019. “Network Curvature as a Hallmark of Brain Structural Connectivity.” Nature Communications 10: 4937. 10.1038/s41467-019-12915-x.

Fischl, Bruce. 2012. “FreeSurfer.” NeuroImage 62 (2): 774–81. 10.1016/j.neuroimage.2012.01.021.

Flamary, Rémi, Nicolas Courty, Alexandre Gramfort, Mokhtar Z. Alaya, Aurélie Boisbunon, Stanislas Chambon, Laetitia Chapel, et al. 2021. “POT: Python Optimal Transport.” Journal of Machine Learning Research 22 (78): 1–8.

Geerligs, Linda, Remco J. Renken, Emi Saliasi, Natasha M. Maurits, and Monicque M. Lorist. 2015. “A Brain-Wide Study of Age-Related Changes in Functional Connectivity.” Cerebral Cortex 25 (7): 1987–99. 10.1093/cercor/bhu012.

Gong, Gaolang, Pedro Rosa-Neto, Felix Carbonell, Zhang J. Chen, Yong He, and Alan C. Evans. 2009. “Age- and Gender-Related Differences in the Cortical Anatomical Network.” Journal of Neuroscience 29 (50): 15684–93. 10.1523/JNEUROSCI.2308-09.2009.

Heuvel, Martijn P. van den, Siemon C. de Lange, Andrew Zalesky, Caio Seguin, B. T. Thomas Yeo, and Ruben Schmidt. 2017. “Proportional Thresholding in Resting-State fMRI Functional Connectivity Networks and Consequences for Patient-Control Connectome Studies: Issues and Recommendations.” NeuroImage 152: 437–49. 10.1016/j.neuroimage.2017.02.005.

Isensee, Fabian, Marianne Schell, Irada Pflueger, Gianluca Brugnara, David Bonekamp, Ulf Neuberger, Antje Wick, et al. 2019. “Automated Brain Extraction of Multisequence MRI Using Artificial Neural Networks.” Human Brain Mapping 40 (17): 4952–64. 10.1002/hbm.24750.

Jeurissen, Ben, Jacques-Donald Tournier, Thijs Dhollander, Alan Connelly, and Jan Sijbers. 2014. “Multi-Tissue Constrained Spherical Deconvolution for Improved Analysis of Multi-Shell Diffusion MRI Data.” NeuroImage 103: 411–26.

Jost, Jürgen, and Shiping Liu. 2014. “Ollivier’s Ricci Curvature, Local Clustering and Curvature-Dimension Inequalities on Graphs.” Discrete and Computational Geometry 51 (2): 300–322. 10.1007/s00454-013-9558-1.

Kellner, Elias, Bibek Dhital, Valerij G. Kiselev, and Marco Reisert. 2016. “Gibbs-Ringing Artifact Removal Based on Local Subvoxel-Shifts.” Magnetic Resonance in Medicine 76 (5): 1574–81.

Koen, Joshua D., and Michael D. Rugg. 2019. “Neural Dedifferentiation in the Aging Brain.” Trends in Cognitive Sciences 23 (7): 547–59. 10.1016/j.tics.2019.04.012.

Maier-Hein, Klaus H., Peter F. Neher, Jean-Christophe Houde, et al. 2017. “The Challenge of Mapping the Human Connectome Based on Diffusion Tractography.” Nature Communications 8: 1349. 10.1038/s41467-017-01285-x.

Malagurski, Brigitta, Franziskus Liem, Jessica Oschwald, Susan Mérillat, and Lutz Jäncke. 2020. “Functional Dedifferentiation of Associative Resting State Networks in Older Adults: A Longitudinal Study.” NeuroImage 214: 116680. 10.1016/j.neuroimage.2020.116680.

Margulies, Daniel S., Satrajit S. Ghosh, Alexandros Goulas, Marcel Falkiewicz, Julia M. Huntenburg, Georg Langs, Gleb Bezgin, et al. 2016. “Situating the Default-Mode Network Along a Principal Gradient of Macroscale Cortical Organization.” Proceedings of the National Academy of Sciences 113 (44): 12574–79. 10.1073/pnas.1608282113.

Markello, Ross D., and Bratislav Mišić. 2021. “Comparing Spatial Null Models for Brain Maps.” NeuroImage 236: 118052. 10.1016/j.neuroimage.2021.118052.

Ollivier, Yann. 2009. “Ricci Curvature of Markov Chains on Metric Spaces.” Journal of Functional Analysis 256 (3): 810–64. 10.1016/j.jfa.2008.11.001.

Raffelt, David, Thijs Dhollander, Jacques-Donald Tournier, Rami Tabbara, Robert E. Smith, Eric Pierre, and Alan Connelly. 2017. “Bias Field Correction and Intensity Normalisation for Quantitative Analysis of Apparent Fibre Density.” In Proceedings of the International Society for Magnetic Resonance in Medicine, 25:3541.

Reese, Timothy G., O. Heid, Robert M. Weisskoff, and Van J. Wedeen. 2003. “Reduction of Eddy-Current-Induced Distortion in Diffusion MRI Using a Twice-Refocused Spin Echo.” Magnetic Resonance in Medicine 49 (1): 177–82.

Roberts, James A., Alistair Perry, Gloria Roberts, Philip B. Mitchell, and Michael Breakspear. 2017. “Consistency-Based Thresholding of the Human Connectome.” NeuroImage 145: 118–29. 10.1016/j.neuroimage.2016.09.053.

Robinson, Tyler D., Jordan A. Chad, Yutong L. Sun, Paul T. H. Chang, and J. Jean Chen. 2026. “Testing Retrogenesis and Physiological Explanations for Tract-Wise White Matter Aging: Links to Developmental Order, Fiber Calibre, and Vascularization.” GeroScience 48 (2): 2401–22. 10.1007/s11357-025-01773-9.

Schaefer, Alexander, Ru Kong, Evan M. Gordon, Timothy O. Laumann, Xi-Nian Zuo, Avram J. Holmes, Simon B. Eickhoff, and B. T. Thomas Yeo. 2018. “Local-Global Parcellation of the Human Cerebral Cortex from Intrinsic Functional Connectivity MRI.” Cerebral Cortex 28 (9): 3095–3114. 10.1093/cercor/bhx179.

Schilling, Kurt G., Justin Blaber, Colin Hansen, Leon Cai, Baxter Rogers, Adam W. Anderson, Seth Smith, et al. 2020. “Distortion Correction of Diffusion Weighted MRI Without Reverse Phase-Encoding Scans or Field-Maps.” PLoS ONE 15 (7): e0236418. 10.1371/journal.pone.0236418.

Schilling, Kurt G., Vishwesh Nath, Colin Hansen, Prasanna Parvathaneni, Justin Blaber, Yurui Gao, Peter Neher, et al. 2019. “Limits to Anatomical Accuracy of Diffusion Tractography Using Modern Approaches.” NeuroImage 185: 1–11. 10.1016/j.neuroimage.2018.10.029.

Seraji, Masoud, Sarah Shultz, Qiang Li, Zening Fu, and Vince D. Calhoun. 2026. “Comparing Nonlinear Trajectories Across Brain Networks: A Key to Understanding Complex Brain Dynamics.” bioRxiv. 10.64898/2026.07.24.740420.

Shafto, Meredith A., Lorraine K. Tyler, Marie Dixon, Jason R. Taylor, James B. Rowe, Rhodri Cusack, Andrew J. Calder, et al. 2014. “The Cambridge Centre for Ageing and Neuroscience (Cam-CAN) Study Protocol: A Cross-Sectional, Lifespan, Multidisciplinary Examination of Healthy Cognitive Ageing.” BMC Neurology 14: 204. 10.1186/s12883-014-0204-1.

Simhal, Anish K., Kimberly L. H. Carpenter, Saad Nadeem, Joanne Kurtzberg, Allen Song, Allen Tannenbaum, Guillermo Sapiro, and Geraldine Dawson. 2020. “Measuring Robustness of Brain Networks in Autism Spectrum Disorder with Ricci Curvature.” Scientific Reports 10: 10819. 10.1038/s41598-020-67474-9.

Smith, Robert E., Jacques-Donald Tournier, Fernando Calamante, and Alan Connelly. 2012. “Anatomically-Constrained Tractography: Improved Diffusion MRI Streamlines Tractography Through Effective Use of Anatomical Information.” NeuroImage 62 (3): 1924–38.

Smith, Robert E., Jacques-Donald Tournier, Fernando Calamante, and Alan Connelly. 2015. “SIFT2: Enabling Dense Quantitative Assessment of Brain White Matter Connectivity Using Streamlines Tractography.” NeuroImage 119: 338–51. 10.1016/j.neuroimage.2015.06.092.

Taylor, Jason R., Nitin Williams, Rhodri Cusack, Tibor Auer, Meredith A. Shafto, Marie Dixon, Lorraine K. Tyler, Cam-CAN, and Richard N. Henson. 2017. “The Cambridge Centre for Ageing and Neuroscience (Cam-CAN) Data Repository: Structural and Functional MRI, MEG, and Cognitive Data from a Cross-Sectional Adult Lifespan Sample.” NeuroImage 144: 262–69. 10.1016/j.neuroimage.2015.09.018.

Touroutoglou, Alexandra, Jiahe Zhang, Joseph M. Andreano, Bradford C. Dickerson, and Lisa Feldman Barrett. 2018. “Dissociable Effects of Aging on Salience Subnetwork Connectivity Mediate Age-Related Changes in Executive Function and Affect.” Frontiers in Aging Neuroscience 10: 410. 10.3389/fnagi.2018.00410.

Tustison, Nicholas J., Brian B. Avants, Philip A. Cook, Yuanjie Zheng, Alexander Egan, Paul A. Yushkevich, and James C. Gee. 2010. “N4ITK: Improved N3 Bias Correction.” IEEE Transactions on Medical Imaging 29 (6): 1310–20. 10.1109/TMI.2010.2046908.

Váša, František, Jakob Seidlitz, Rafael Romero-Garcia, Kirstie J. Whitaker, Gideon Rosenthal, Petra E. Vértes, Maxwell Shinn, et al. 2018. “Adolescent Tuning of Association Cortex in Human Structural Brain Networks.” Cerebral Cortex 28 (1): 281–94. 10.1093/cercor/bhx249.

Veraart, Jelle, Dmitry S. Novikov, Daan Christiaens, Benjamin Ades-Aron, Jan Sijbers, and Els Fieremans. 2016. “Denoising of Diffusion MRI Using Random Matrix Theory.” NeuroImage 142: 394–406.

Vos de Wael, Reinder, Oualid Benkarim, Casey Paquola, Sara Lariviere, Jessica Royer, Shahin Tavakol, Ting Xu, et al. 2020. “BrainSpace: A Toolbox for the Analysis of Macroscale Gradients in Neuroimaging and Connectomics Datasets.” Communications Biology 3: 103. 10.1038/s42003-020-0794-7.

West, Robert L. 1996. “An Application of Prefrontal Cortex Function Theory to Cognitive Aging.” Psychological Bulletin 120 (2): 272–92. 10.1037/0033-2909.120.2.272.

Yadav, Yasharth, Pavithra Elumalai, Nitin Williams, Jürgen Jost, and Areejit Samal. 2023. “Discrete Ricci Curvatures Capture Age-Related Changes in Human Brain Functional Connectivity Networks.” Frontiers in Aging Neuroscience 15: 1120846. 10.3389/fnagi.2023.1120846.

Yeo, B. T. Thomas, Fenna M. Krienen, Jorge Sepulcre, Mert R. Sabuncu, Danial Lashkari, Marisa Hollinshead, Joshua L. Roffman, et al. 2011. “The Organization of the Human Cerebral Cortex Estimated by Intrinsic Functional Connectivity.” Journal of Neurophysiology 106 (3): 1125–65.

Zhou, Zhen, Bruce Fischl, and Iman Aganj. 2025. “Harmonization of Structural Brain Connectivity Through Distribution Matching.” Human Brain Mapping 46 (9): e70257. 10.1002/hbm.70257.

Zhu, Silei, Nicola K. Dinsdale, Saad Jbabdi, Karla L. Miller, and Amy F. D. Howard. 2026. “Microscopy-Informed Structural Connectivity Mapping in the in Vivo Human Brain via Domain Adaptation.” bioRxiv. 10.64898/2026.06.14.732211.

