## Supplementary Material for "Ageing conserves and redistributes local geometry in the human structural connectome: an Ollivier-Ricci curvature analysis across the adult lifespan"

Supplementary Materials

This document contains two annexes, each self-contained.

**Annex A** derives the quantities used in the main text, states the results that motivate their use, and documents the numerical validation of the implementation. It is written to be readable by someone who has not previously met optimal transport or discrete curvature.

**Annex B** reports analyses that were run and are not required by the argument of the main text, either because they returned nothing, because they support a claim already made by a shorter route, or because they are descriptive rather than inferential. Several of them constrain the interpretation, and two are informative precisely because they are null.

**Annex C** gives the processing chain in full, from raw diffusion data to the weighted connectivity matrix, naming every piece of software, every command-line option and the estimator behind each step, so that the reconstruction can be repeated exactly. The Methods section of the manuscript describes the same chain at the level a reader needs to follow the argument.

#### Figures

B1 The increasing and the decreasing node sets both follow nonlinear age courses, and differ in amplitude rather than in shape . . . . . . . . . . 13
B2 The age effect on curvature is present within every stratum of edge length . . . . . . . . 15
B3 The concentration of decreasing nodes in the default mode network appears for curvature alone . . . . . . . . . . 17
B4 Curvature exceeds what the degree sequence alone would produce, and the null carries its own age trend . . . . . . . . . . 18
B5 The age effect does not depend on the SIFT2 reweighting . . . . . . . . . . 19
B6 The shape of the age course is preserved across covariate specifications; only the variance attributed to age changes . . . . . . . . . . 20
B7 Cohort composition and the quantities motivating harmonisation . . . . . . . . . . 22
B8 Curvature does not outperform magnitude at predicting age . . . . . . . . . . 22

B9 The decreasing nodes are not distinguishable by classical nodal properties . . . . . . . . 23
C1 Anatomically constrained tractography in one participant . . . . . . . . . . . . . . . . . . 28
C2 Connectivity matrices at three levels . . . . . . . . . . . . . . . . . . . . . . . . . . . . . . . . 30

### Tables

A1 Spatial smoothness of the observed effect map and of the two null models . . . . . . . . 8
A2 Numerical validation of the curvature implementation . . . . . . . . . . . . . . . . . . . . 10
B1 Age effect on curvature within each stratum of edge length . . . . . . . . . . . . . . . . . 14
B2 Specificity of the network enrichment across four nodal measures . . . . . . . . . . . . . 16
B3 Age effect on mean curvature under three covariate specifications . . . . . . . . . . . . . 19
B4 All modelled network blocks . . . . . . . . . . . . . . . . . . . . . . . . . . . . . . . . . . . 21
C1 Diffusion MRI acquisition parameters . . . . . . . . . . . . . . . . . . . . . . . . . . . . . . 32
C2 Processing environment . . . . . . . . . . . . . . . . . . . . . . . . . . . . . . . . . . . . . . 32

### Annex A. Mathematical background and validation

This annex derives the quantities used in the main text, states the results that motivate their use, and documents the numerical validation of the implementation. It is written to be self-contained for a reader who has not met optimal transport or discrete curvature before.

---

##### A.0 Supporting acquisition detail

The main text reports the diffusion and T1-weighted parameters. Two further sequences were acquired in the same session and bear on preprocessing rather than on the analysis, and are recorded here for completeness.

A T2-weighted SPACE image was acquired at 1.0 mm isotropic resolution, 192 slices per slab, field of view 256 mm, base resolution 256, TR 2,800 ms, TE 408 ms, GRAPPA factor 2, acquisition time 4 min 30 s. A dual-echo gradient-echo field map was acquired at 3.0 by 3.0 by 3.7 mm over 32 slices, field of view 192 mm, TR 400 ms, echo times 5.19 and 7.65 ms, acquisition time 54 s, and supports correction of susceptibility-induced distortion in the diffusion volumes.

Gradient nonlinearity was corrected using the vendor gradient coefficient file distributed with the Cam-CAN release.

##### A.1 From Riemannian to discrete curvature

In Riemannian geometry, Ricci curvature answers a local question: do two geodesics that leave nearby points in parallel directions converge, stay parallel, or diverge? On a sphere they converge, on a plane they stay parallel, on a saddle they diverge. Equivalently, curvature controls how the volume of a small ball compares with the Euclidean volume of the same radius.

A graph has neither smooth geodesics nor a metric tensor, so this definition does not transfer. Ollivier (2009) restated the question in probabilistic terms, in a form that survives discretisation:

> Is the neighbourhood of $x$ closer to the neighbourhood of $y$ than $x$ is to $y$?

If neighbourhoods approach each other faster than their centres, the space closes, which is positive curvature. If they separate faster, the space opens, which is negative curvature. Making this operational requires a probability measure at each node and a distance between measures.

###### A.1.1 The lazy random walk

For a weighted graph with affinities $w_{xy} > 0$, the one-step lazy random-walk measure at $x$ is

$$m_x^\alpha(z) = \begin{cases} \alpha, & z = x, \\ (1-\alpha)\,\dfrac{w_{xz}}{\sum_{k\sim x} w_{xk}}, & z \sim x, \\ 0, & \text{otherwise.} \end{cases}$$

The laziness $\alpha$ is the probability of remaining in place. It exists for a technical reason: on a bipartite graph a non-lazy walk oscillates and never converges, and laziness removes the oscillation. Its practical effect is to rescale curvature without reordering edges, which we verify empirically in the main text.

The measure has $\deg(x) + 1$ points of support and sums to one by construction.

###### A.1.2 Wasserstein-1 distance

Given two probability measures $\mu$ and $\nu$ on the node set and a ground cost $d(a, b)$, the Wasserstein-1 distance is

$$W_1(\mu, \nu) = \min_{\pi \in \Pi(\mu,\nu)} \sum_{a,b} \pi(a, b)\, d(a, b),$$

where $\Pi(\mu, \nu)$ is the set of couplings, that is non-negative matrices whose row sums reproduce $\mu$ and whose column sums reproduce $\nu$. The minimiser $\pi^\star$ is the optimal transport plan, and the value is the least total cost of rearranging one measure into the other. This is a linear programme in $\pi$, and for the small supports involved here, at most a few hundred points, it is solved exactly in microseconds.

The ground cost is the shortest-path distance computed on the edge distances $d_{ij}$. Note that $W_1$ therefore sees structure beyond the immediate neighbourhood, since a shortest path may traverse several edges.

###### A.1.3 The curvature

$$\boxed{\kappa(x, y) = 1 - \frac{W_1\big(m_x^\alpha,\, m_y^\alpha\big)}{d(x, y)}}$$

Read as a comparison of two costs. The denominator $d(x, y)$ is the cost of dragging the mass along the edge itself, with no shortcut. The numerator is the cost optimal transport actually pays. Their ratio is one when no shortcut exists, less than one when alternative short routes are available, and greater than one when the neighbourhood of $y$ lies further from that of $x$ than the centres lie from each other.

Hence positive curvature means local redundancy and negative curvature means a bottleneck. Since $W_1 \geq 0$, we have $\kappa \leq 1$ always.

A point worth making explicit, because it is where the intuition of "bottleneck" becomes quantitative: mass crossing a bridge must not only traverse it but then redistribute on the far side. It is this second cost that pushes $W_1$ above $d(x, y)$ and drives curvature negative. A bottleneck is expensive not because it is narrow but because everything passing through it must then spread out.

---

##### A.2 Closed forms used for validation

Three families admit exact solutions, and they were used to validate the implementation before it touched data.

###### A.2.1 Complete graph $K_n$

All edges have unit affinity and unit distance, and every pair of nodes is adjacent, so $d(a, b) = 1$ for $a \neq b$. Consider the edge $(x, y)$. The measures $m_x$ and $m_y$ agree on the $n - 2$ nodes adjacent to both, each carrying $(1 - \alpha)/(n - 1)$. They differ only in that $m_x$ places $\alpha$ at $x$ and $(1 - \alpha)/(n - 1)$ at $y$, while $m_y$ does the reverse.

The shared mass costs nothing to move. The mass that must move is $\alpha - (1 - \alpha)/(n - 1)$, travelling a distance of one. Therefore

$$W_1 = \alpha - \frac{1-\alpha}{n-1}, \qquad \kappa = 1 - \alpha + \frac{1-\alpha}{n-1}.$$

At $\alpha = 0.5$ and $n = 8$ this gives $\kappa = 0.5714$, and the implementation returns the same to sixteen decimal places.

###### A.2.2 Cycle $C_n$ with $n \geq 6$

Every node has degree two. The neighbourhoods of adjacent nodes $x$ and $y$ are disjoint apart from $x$ and $y$ themselves, and for $n \geq 6$ the cycle is locally a path, so no shortcut exists. The transport cost equals the edge length exactly and

$$\kappa = 0.$$

The cycle is the discrete flat case, and it is a useful check precisely because it must return exactly zero rather than approximately zero.

###### A.2.3 Infinite $d$-regular tree

A tree has no cycles, so no alternative routes exist at all. Mass must not only cross the edge but also fan out into $d-1$ new branches on the far side. The result is

$$\kappa = -\frac{2(1-\alpha)(d-2)}{d}.$$

For $d = 2$ the tree is a path and curvature vanishes, consistent with the cycle. Curvature becomes more negative as $d$ grows, since more mass must fan out.

**A finite-sample caveat that matters in practice.** Random $d$-regular graphs are only locally tree-like, and the short cycles that remain add exactly the redundancy that raises curvature. Measured values therefore sit systematically above the infinite-tree prediction, and the gap widens with $d$: at $n = 400$ we obtained $-0.325$ against $-0.333$ for $d = 3$, and $-0.522$ against $-0.600$ for $d = 5$. This is not an implementation error, and a validation suite that demanded exact agreement here would be wrong to do so.

---

##### A.3 What curvature controls

Three results justify preferring curvature over an index defined by convenience.

**Mixing.** If curvature is bounded below by $\kappa_{\min}$, the spectral gap $\lambda$ of the associated Markov chain satisfies $\lambda \geq \kappa_{\min}$. Since mixing time scales as $1/\lambda$, positive curvature guarantees fast diffusion toward equilibrium. The bound is on the minimum, not the mean, a distinction worth respecting: summaries based on the mean are descriptively useful but are not what the theorem constrains.

**Perturbation response.** Through the fluctuation theorem of Demetrius, network entropy relates positively to the rate at which a system returns to its state after perturbation, and Ollivier-Ricci curvature serves as a proxy for that entropy (Sandhu et al., 2016). The resulting statement is one of co-monotonicity, $\Delta\kappa \times \Delta(\text{robustness}) \geq 0$, rather than an equality.

**Message passing.** Edges of strongly negative curvature are the bottlenecks at which information is squeezed in graph neural networks, which is the phenomenon of over-squashing (Topping et al.,

2022). This is not used in the present study but explains why curvature-guided rewiring has become a standard preprocessing step in geometric deep learning. Discrete Ricci flow on the same quantity underlies curvature-based community detection (Ni et al., 2019), and the first application of Ollivier-Ricci curvature to structural connectomes established its sensitivity to age and to clinical group (Farooq et al., 2019).

---

##### A.4 The affinity-to-distance transform

Curvature requires a cost, whereas a connectome weight is an affinity: a large $w$ means a strong connection, which should correspond to a *short* distance. Two monotone decreasing transforms are natural.

The inverse, $d = 1/w$, is bounded away from zero for finite weights. The logarithm, $d = -\log(w/w_{\max})$, is additive along paths, which is desirable for shortest-path quantities, but sends the strongest edge to $d = 0$. Since $\kappa = 1 - W_1/d$ divides by $d$, this is a pole, and curvature diverges. Applied to these data, the logarithmic transform produced mean curvatures on the order of $-10^5$.

We therefore used $d = 1/w$ throughout. The library shifts the logarithmic variant by one and emits a warning when the ratio of largest to smallest distance exceeds $10^8$, but the inverse is the appropriate default for curvature.

The direction of the transform is not a matter of taste. Passing raw affinities where distances are expected inverts the sign of the result on a substantial fraction of edges. In these data, the two conventions produced nodal maps correlated at $\rho = -0.29$, with 43% of edges changing sign, and neither map looks obviously wrong on inspection.

---

##### A.5 The threshold-depth confound

This is a general problem with proportional thresholding, not specific to curvature, and it is worth stating in its own terms.

Let participant $i$ have $P_i$ positive edges and let the threshold retain $E$ edges for everyone. The retained set is the strongest $E$ of $P_i$, so the retained *fraction* is $E/P_i$, which varies across participants whenever $P_i$ does. A participant with many positive edges contributes a highly selective skeleton of strong connections; a participant with few contributes nearly their whole graph, weak edges included.

The two graphs have the same size but not the same composition. If $P_i$ covaries with a variable of interest, so does composition, and any association with that variable is confounded. In these data $P_i$ ranged from 4,116 to 16,253, so the retained fraction ranged from 0.72 to 0.18, and $P_i$ declined with age.

**The control.** Subsample the positive edges of every participant uniformly to a common supply $P_{\min}$ before thresholding. The retained fraction becomes $E/P_{\min}$ for everyone. Sampling must ignore weight: sampling proportional to weight would reintroduce exactly the selectivity the control is meant to remove.

**The correction that the control requires.** Subsampling discards information and therefore attenuates any correlation on its own, independently of whether a confound existed. Without correcting for this, the control always appears to have destroyed part of the effect. The reliability of the subsampled measure is the correlation between two independent subsamples of the same participants, $r_{xx}$, and the disattenuated estimate is

$$\rho_{\text{corrected}} = \frac{\rho_{\text{observed}}}{\sqrt{r_{xx}}}.$$

---

#### A.6 Null models

##### A.6.1 Degree-preserving rewiring

Two edges $(a, b)$ and $(c, d)$ with four distinct endpoints are replaced by $(a, d)$ and $(c, b)$, provided neither replacement already exists. Every node degree is preserved exactly, and weights travel with the edges so the weight distribution is preserved exactly as well. Swaps that would disconnect the graph are rejected.

The normalised statistic

$$z_i = \frac{\bar{\kappa}_i^{\text{obs}} - \langle \bar{\kappa}_i^{\text{null}} \rangle}{\text{sd}(\bar{\kappa}_i^{\text{null}})}$$

answers the question a reviewer will ask: how much curvature does this brain carry beyond what its degree sequence alone would produce?

An implementation note, since it determines whether the analysis is feasible at all. Maintaining the edge list as an array with an adjacency matrix for membership tests makes each swap $O(1)$; rebuilding the edge list per swap makes it $O(E)$ and takes minutes per participant instead of milliseconds.

##### A.6.2 Spatial rotation

Cortical maps are spatially autocorrelated, so permuting parcel labels destroys that structure and yields a null that is far too liberal. The rotation null instead rotates parcel centroids on the sphere by a random orthogonal transformation and reassigns each parcel to its nearest rotated neighbour, preserving the spatial autocorrelation and destroying only the alignment with anatomy (Alexander-Bloch et al., 2018).

We used the one-to-one reassignment of Váša et al. (2018), in which each target parcel is matched exactly once, so that each null map is a strict permutation of the observed map and the marginal distribution is identical. The alternative nearest-neighbour rule permits duplicates and omissions, which perturbs the marginal slightly. For 200 parcels the cost of the one-to-one rule is negligible.

The sign of an eigenvector is arbitrary, and the same is true of any orientation convention used to build the initial correspondence; both orientations must be tried and the better retained. Failing to do so makes the spectral initialisation worse than no initialisation at all.

**Diagnostic.** A null model should be verified rather than assumed, and the verification is cheap. Correlating each parcel value with the mean of its six nearest neighbours on the sphere gives a smoothness index, which a valid rotation null must retain.

**Table A1:** Smoothness of the observed nodal effect map compared with rotation nulls and with naive label permutation, measured as the correlation between each parcel and the mean of its six nearest neighbours on the sphere.

| map | smoothness |
|---|---|
| observed nodal effect map | 0.97 |
| rotation nulls | 0.72, SD 0.06 |
| naive label permutation | 0.00, SD 0.09 |

The rotation nulls sit below the observed map, as they must, since rotation displaces each value from its own neighbourhood. They sit far above naive permutation, which is the point: a null scoring near zero has destroyed the autocorrelation it was meant to preserve, and would indicate a failure of the centroid construction rather than a conservative test.

---

##### A.7 Testing a difference in shape

The dissociation between two age courses is not a comparison of two correlations. Two measures can both change with age, at different rates, and still share the same *shape*; the claim of a dissociation is a claim about shape.

Standardise both measures within the cohort and take the within-participant difference

$$D_i = z(\bar{\kappa}_i) - z(s_i)\,.$$

Under the null hypothesis that the two follow the same age course up to scale and sign, $\mathbb{E}[D \mid \text{age}]$ is constant. Testing the dissociation therefore reduces to testing whether $D$ depends on age, which is an ordinary $F$ test of the spline block against a model with no age term.

This has three advantages over an interaction term in long format. The pairing is exact and requires no random-effects model. The assumptions are those of ordinary least squares and can be checked directly. And the fitted curve of $D$ is interpretable: it shows *where* in the lifespan the two clocks diverge, not merely that they do.

A distribution-free counterpart flips the sign of each participant's difference, which is exchangeable under the same null, and requires no normality assumption.

**Effect size.** Report the age at which half of the total variation of each fitted curve has occurred,

$$a_{1/2} = \min\left\{ a : \int_{a_{\min}}^{a} |f'(u)|\, du \geq \tfrac{1}{2} \int_{a_{\min}}^{a_{\max}} |f'(u)|\, du \right\},$$

computed on the fitted spline. The difference between the two half-change ages, with a bootstrap interval over participants, expresses the dissociation in years rather than in units of variance.

---

##### A.8 Breaking circularity

When a set of nodes is defined by a statistic and its composition is then tested with the same statistic on the same data, the test is circular and its $p$ value is uninterpretable. Three routes break it, and they fail independently, which is why all three are reported.

**Remove the selection.** Contrast the mean nodal statistic inside and outside a network with no selection, no threshold and no correction. Nothing is chosen, so nothing can be chosen circularly. This is the most conservative route and will miss effects that live in the tail rather than in the mean.

**Separate selection from inference.** Define the set in one half of the participants and evaluate it in the held-out half. Selection and inference then use different data, and the circularity is removed by construction rather than by argument. Repeating over many splits gives a distribution rather than a single number.

**Match the procedure.** Apply the identical selection and null procedure to alternative statistics, with the set size fixed at the value observed for the statistic of interest. If the result appears for all of them, it is a property of the data rather than of the statistic; if it appears for one, that is specificity. This route tests specificity but not circularity, which is why it is insufficient on its own.

---

##### A.9 Suppression, and why an adjustment can raise an estimate

A covariate that lies on the causal path between predictor and outcome is a mediator; adjusting for it estimates a direct rather than a total effect, and the adjusted estimate is normally smaller.

A covariate can also act as a *suppressor*. If $C$ correlates with the predictor $A$ and with the outcome $Y$ in directions such that its indirect contribution partially cancels the direct one, then removing $C$ from the residual variance increases the apparent association between $A$ and $Y$. Formally, for standardised variables the partial correlation

$$r_{AY \cdot C} = \frac{r_{AY} - r_{AC} r_{YC}}{\sqrt{\left(1 - r_{AC}^2\right)\left(1 - r_{YC}^2\right)}}$$

exceeds $r_{AY}$ whenever $r_{AC} r_{YC}$ carries the sign opposite to $r_{AY}$.

The practical consequence is that an unadjusted estimate in the presence of a suppressor is conservative rather than inflated. This is the opposite of the usual reading of a confound, and it should be stated explicitly when it occurs, since a reader who sees an estimate rise after adjustment will otherwise suspect an error.

---

##### A.10 Numerical validation summary

**Table A2:** Numerical validation of the curvature implementation against closed forms and against the properties each null model is required to preserve.

| Check | Expected | Obtained |
|---|---|---|
| $K_n$, twelve combinations of $n$ and $\alpha$ | closed form | agreement to $2.2 \times 10^{-16}$ |
| $C_n$, $n \in \{8, 12, 20\}$ | exactly 0 | 0 to machine precision |
| Random $d$-regular, $n = 400$ | above the infinite-tree value | confirmed, gap widening with $d$ |
| Laziness $\alpha$ from 0.0 to 0.9 | monotone rescaling | edge ranking preserved, $\rho \geq 0.90$ |
| Rotation nulls, marginal distribution | identical to observed | exact, per hemisphere |
| Rotation nulls, spatial smoothness | high but below observed | 0.72 against 0.97; naive permutation 0.[illegible] |
| Degree-preserving rewiring | degrees and weights exactly preserved | confirmed for all participants |

---

##### References for this annex

Alexander-Bloch, A. F., Shou, H., Liu, S., Satterthwaite, T. D., Glahn, D. C., Shinohara, R. T., Vandekar, S. N., & Raznahan, A. (2018). On testing for spatial correspondence between maps of human brain structure and function. *NeuroImage*, 178, 540-551.

Farooq, H., Chen, Y., Georgiou, T. T., Tannenbaum, A., & Lenglet, C. (2019). Network curvature as a hallmark of brain structural connectivity. *Nature Communications*, 10, 4937.

Fischl, B. (2012). FreeSurfer. *NeuroImage*, 62(2), 774-781.

Flamary, R., Courty, N., Gramfort, A., et al. (2021). POT: Python Optimal Transport. *Journal of Machine Learning Research*, 22(78), 1-8.

Margulies, D. S., Ghosh, S. S., Goulas, A., et al. (2016). Situating the default-mode network along a principal gradient of macroscale cortical organization. *PNAS*, 113(44), 12574-12579.

Ni, C.-C., Lin, Y.-Y., Luo, F., & Gao, J. (2019). Community detection on networks with Ricci flow. *Scientific Reports*, 9, 9984.

Ollivier, Y. (2009). Ricci curvature of Markov chains on metric spaces. *Journal of Functional Analysis*, 256(3), 810-864.

Sandhu, R. S., Georgiou, T. T., & Tannenbaum, A. R. (2016). Ricci curvature: An economic indicator for market fragility and systemic risk. *Science Advances*, 2(5), e1501495.

Schaefer, A., Kong, R., Gordon, E. M., et al. (2018). Local-global parcellation of the human cerebral cortex from intrinsic functional connectivity MRI. *Cerebral Cortex*, 28(9), 3095-3114.

Smith, R. E., Tournier, J.-D., Calamante, F., & Connelly, A. (2015). SIFT2: Enabling dense quantitative assessment of brain white matter connectivity using streamlines tractography. *NeuroImage*, 119, 338-351.

Topping, J., Di Giovanni, F., Chamberlain, B. P., Dong, X., & Bronstein, M. M. (2022). Understanding over-squashing and bottlenecks on graphs via curvature. *ICLR*.

Váša, F., Seidlitz, J., Romero-Garcia, R., et al. (2018). Adolescent tuning of association cortex in human structural brain networks. *Cerebral Cortex*, 28(1), 281-294.

Vos de Wael, R., Benkarim, O., Paquola, C., et al. (2020). BrainSpace: a toolbox for the analysis of macroscale gradients in neuroimaging and connectomics datasets. *Communications Biology*, 3, 103.

### Annex B. Extended results

The main text reports what the argument requires. This annex reports analyses that were run and are not needed for the argument, either because they returned nothing, because they support a claim already made by a shorter route, or because they are descriptive rather than inferential. Several of them constrain the interpretation, and two are informative precisely because they are null.

All analyses use the same 307 participants, the same graphs at fixed density 0.15, and the same conventions as the main text.

---

##### B.1 The two node populations have different trajectories, not different shapes

The main text reports that 57 nodes increase in curvature with age and 10 decrease. It does not report what the two groups do as a function of age, which is worth stating because the answer is not the obvious one.

Both populations follow nonlinear age courses. Taking the mean curvature of each group per participant and fitting the same spline model used throughout, the increasing nodes give $R^2 = 0.379$ with $p_{\text{nonlinear}} = 8.7 \times 10^{-8}$, and the decreasing nodes give $R^2 = 0.084$ with $p_{\text{nonlinear}} = 1.1 \times 10^{-4}$. The decreasing set therefore has a genuine and nonlinear age course of its own; what distinguishes it is not the absence of structure but the amount of variance that age explains, which is between four and five times smaller.

Two readings are available and the data do not separate them. The decreasing nodes may be a distinct subsystem with a weaker but real age course, or they may be a heterogeneous set unified only by sign, in which case the residual nonlinearity is inherited from the global trend they oppose. A test that would separate them is whether the decreasing nodes covary with one another across participants more than they covary with the increasing nodes, which is a straightforward analysis on the existing data and is left for future work.

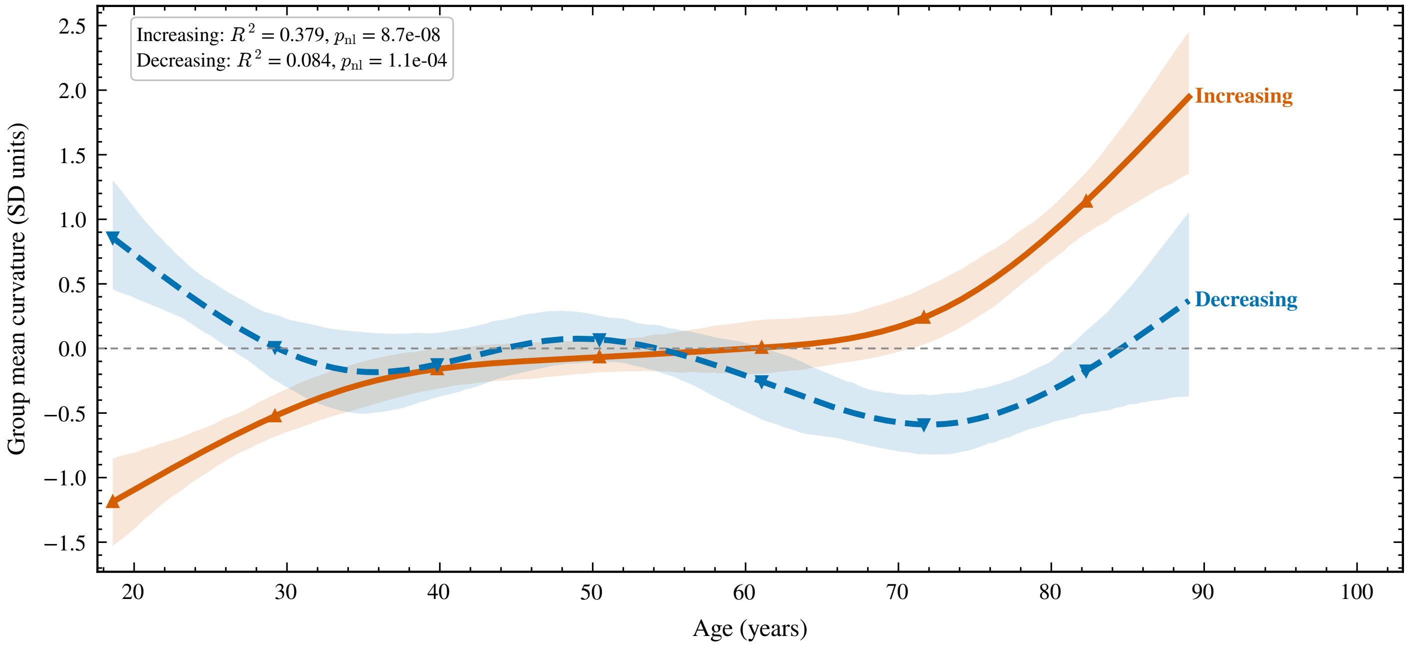

**Figure B1:** The increasing and the decreasing node sets both follow nonlinear age courses, and differ in amplitude rather than in shape. Group mean nodal curvature for each set, expressed in cohort standard-deviation units and adjusted for sex. Shaded bands are bootstrap 95% confidence intervals over 800 resamples. Increasing nodes: $R^2 = 0.379$, $p(\text{nonlinearity}) = 8.7 \times 10^{-8}$. Decreasing nodes: $R^2 = 0.084$, $p(\text{nonlinearity}) = 1.1 \times 10^{-4}$. $n = 307$.

##### B.2 The decreasing nodes are not spatially clustered

Concentration in a network and concentration in space are different claims, and only the first survives. Taking the mean pairwise angle between the decreasing nodes on the sphere as a measure of spatial dispersion, the observed value was 1.127 radians against 0.998 under spatially rotated null maps ($z = 0.63$, $p = 0.564$). The decreasing nodes are, if anything, slightly more dispersed than chance, and certainly not clustered.

This matters for interpretation. A set of nodes concentrated in one cortical territory would invite a focal explanation, such as a locally acting vascular or metabolic factor. A set distributed across the cortex but concentrated in one functional network invites a connectional explanation instead. The result is consistent with the second and not the first, and it also rules out the possibility that the network enrichment reported in the main text is an artefact of the default mode network occupying contiguous cortex.

##### B.3 The nodal effect map is bilaterally symmetric

Pairing each left-hemisphere parcel with the right-hemisphere parcel whose mirrored centroid is nearest, the two members of a pair agreed in the sign of their age effect in 77% of pairs, against 61% under spatially rotated nulls ($z = 3.00$, $p = 8.0 \times 10^{-4}$).

Bilateral symmetry is weak evidence taken alone, because the parcellation is itself approximately symmetric. It becomes informative when combined with B.2: the effect map is not spatially clustered, yet it is mirror-symmetric, which is difficult to produce by any noise process that does not respect the anatomy. We take it as a consistency check on the nodal map rather than as a finding.

##### B.4 The age effect is present within every length stratum

The main text reports the length-stratified analysis in a sentence. The full result is given here because it addresses what is, in our judgement, the most serious threat to the interpretation.

Tractography systematically underestimates long connections, and ageing reduces them. If curvature increased with age only because low-curvature long-range edges were selectively lost, the association should be confined to, or at least strongest in, the long stratum.

**Table B1:** Age effect on mean curvature computed within each tercile of edge length, with edges assigned to strata by their group median length.

| stratum | edges | $\rho$ with age | p | p nonlinear |
|---|---|---|---|---|
| short | 11,151 | +0.354 | $1.7 \times 10^{-10}$ | $4.5 \times 10^{-9}$ |
| medium | 4,440 | +0.371 | $2.0 \times 10^{-11}$ | $7.7 \times 10^{-10}$ |
| long | 4,309 | +0.297 | $1.2 \times 10^{-7}$ | $5.3 \times 10^{-8}$ |

The prediction fails in both respects. The association is present in all three strata with the same sign and comparable magnitude, and it is weakest, not strongest, in the long stratum. Mean length of retained edges did decline with age ($r = -0.310$), and carrying it as a covariate left $R^2 = 0.111$ with $p_{\text{nonlinear}} = 4.9 \times 10^{-7}$.

This does not establish that tractography recovered long connections accurately. It establishes that the age effect on curvature does not depend on the edges where that inaccuracy is concentrated.

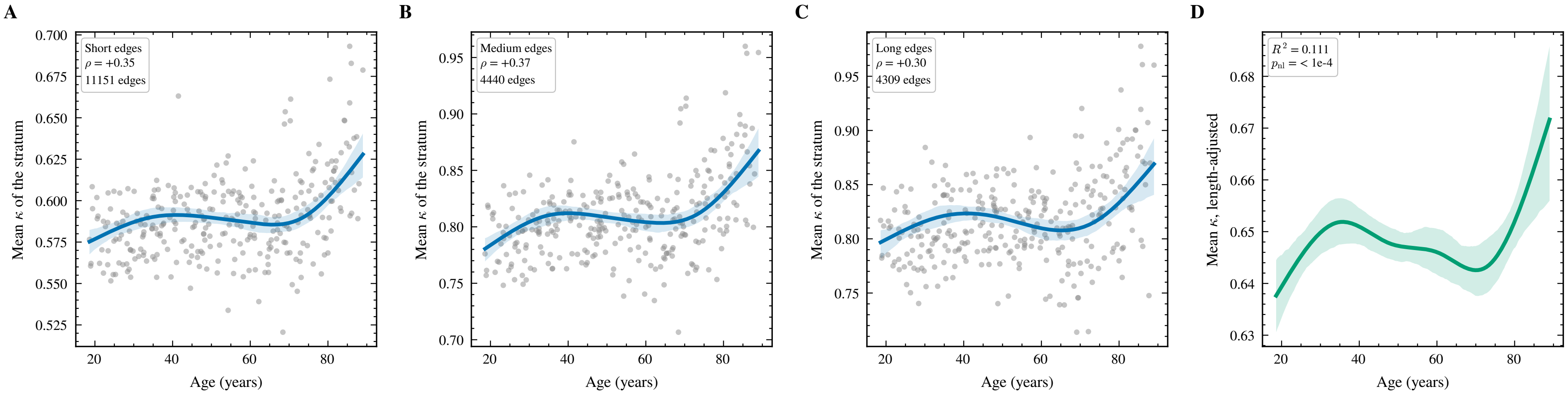

**Figure B2:** The age effect on curvature is present within every stratum of edge length. (A) Mean curvature of the short third of edges against age. (B) Mean curvature of the middle third. (C) Mean curvature of the long third. (D) Mean curvature over all retained edges, with the mean length of those edges carried as an additional covariate. Edges were assigned to terciles by their group median length, so the same edge falls in the same stratum for every participant. A length artefact predicts the effect to be confined to, or strongest in, the long stratum. It is weakest there. Mean retained-edge length declined with age $(r = -0.310)$. $n = 307$.

##### B.5 Specificity across four nodal measures

The main text reports that applying the identical selection and rotation procedure to strength, degree and clustering produced no default mode enrichment. The four tests are given together here, with the set size fixed at 10 nodes throughout so that the comparison is exact.

**Table B2:** Default mode enrichment of the decreasing node set under four nodal measures, with the identical selection and rotation procedure and the set size fixed throughout.

| measure | DMN fraction | expected under rotation | z | p |
|---|---|---|---|---|
| curvature | 0.70 | 0.214 | 3.53 | $1.4 \times 10^{-3}$ |
| strength | 0.30 | 0.246 | 0.37 | 0.73 |
| degree | 0.40 | 0.232 | 1.20 | 0.26 |
| clustering | 0.30 | 0.235 | 0.79 | 0.70 |

Only curvature exceeds the Bonferroni-corrected threshold across the four tests. The three magnitude and topology measures do select node sets, and those sets are not empty, but their network composition is indistinguishable from what spatial rotation produces.

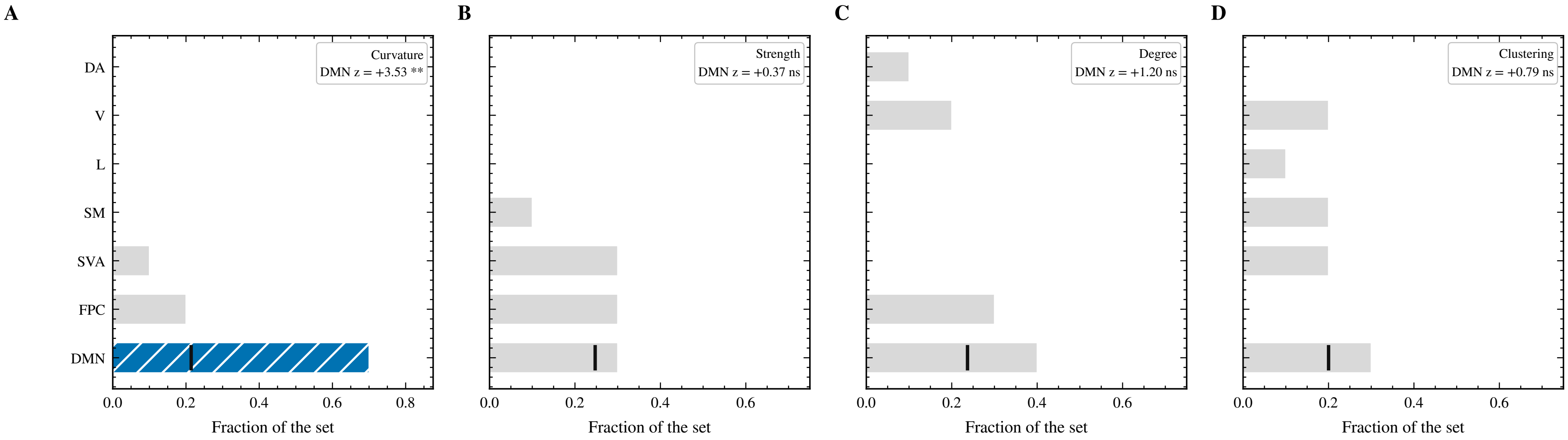

**Figure B3:** The concentration of decreasing nodes in the default mode network appears for curvature alone. (A) Network composition of the ten most strongly decreasing nodes of the curvature age-effect map. (B) The identical selection applied to the nodal strength map. (C) The identical selection applied to the nodal degree map. (D) The identical selection applied to the nodal clustering map. Abbreviations: V, visual network; SM, somatomotor network; DA, dorsal attention network; SVA, salience and ventral attention network; L, limbic network; FPC, frontoparietal control network; DMN, default mode network. Set size fixed at 10 nodes in all four panels so the comparison is exact. The vertical mark gives the default mode proportion expected under 5000 spatial rotations. Only curvature exceeds the Bonferroni-corrected threshold across the four tests. $n = 307$.

##### B.6 The degree-preserving null carries its own age trend

Normalising each participant's mean curvature against five degree-preserving rewirings gave a residual association with age of $\rho = 0.181$ ($p = 1.5 \times 10^{-3}$), which the main text reports. The intermediate quantity is also informative: the mean curvature of the rewired graphs itself increased with age.

This is expected and worth stating explicitly. Degree-preserving rewiring holds the degree sequence fixed, and the degree sequence changes with age; a null built from an ageing brain is itself an ageing null. The normalised statistic therefore answers a sharper question than the raw one. It asks not whether curvature increases with age, which it does, but whether it increases beyond what the degree sequence alone would produce. That it does, at $\rho = 0.181$, is the result; that the null moves too is the reason the normalisation was necessary.

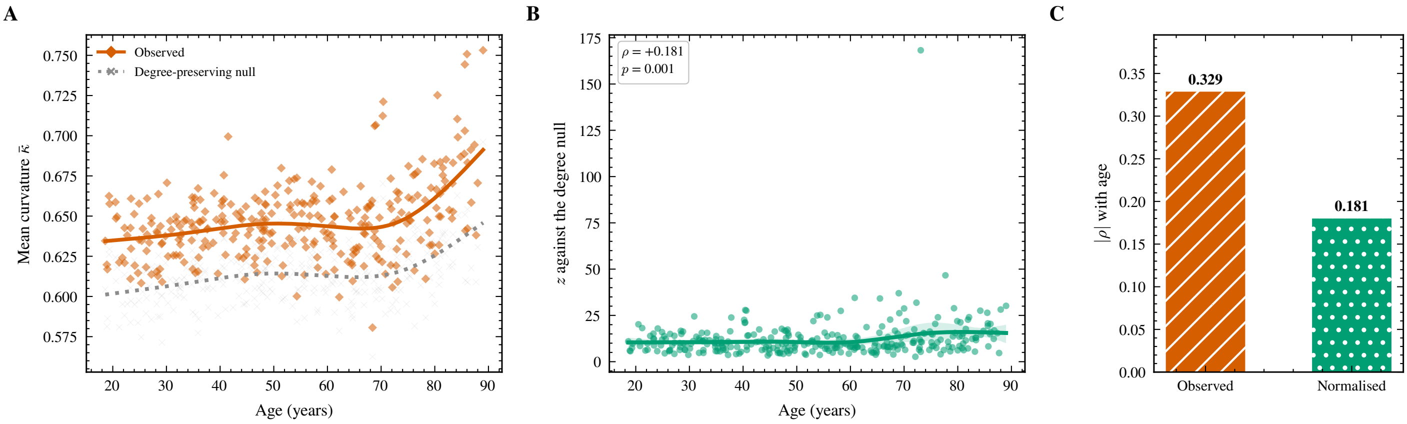

**Figure B4:** Curvature exceeds what the degree sequence alone would produce, and the null carries its own age trend. (A) Observed mean curvature and the mean over five degree-preserving rewirings per participant, against age, with sex-adjusted spline fits. The null rises with age because the degree sequence it preserves does. (B) The normalised statistic, expressing each participant's observed curvature in standard deviations of their own null distribution. (C) Absolute rank correlation with age before and after normalisation. Five rewirings per participant; every node degree preserved exactly and edge weights carried with the edges. $n = 307$.

##### B.7 Replication with unweighted streamline counts

Replacing SIFT2 coefficients with raw streamline counts and repeating the entire pipeline gave $\rho = 0.339$ with age, against 0.329 for SIFT2. The nodal maps of the age effect derived from the two weightings correlated at $\rho = 0.88$.

SIFT2 reweights streamlines according to a diffusion model, and a result that depended on that reweighting would depend on a modelling choice. This one does not. The near-identity of the two estimates also indicates that the age effect is not carried by the small number of edges whose weight the SIFT2 correction changes most.

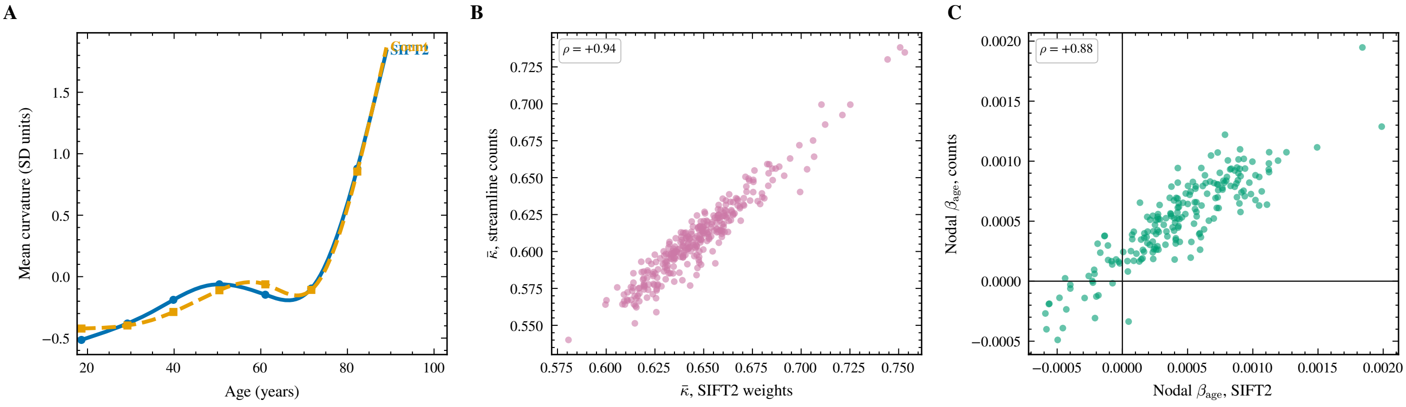

**Figure B5:** The age effect does not depend on the SIFT2 reweighting. (A) Age courses of mean curvature under the two weightings, standardised to a common scale. (B) Participant-level agreement between the two weightings. (C) Agreement of the nodal age-effect maps derived from each. SIFT2 reweights streamlines according to a diffusion model; raw counts do not. A result carried by that reweighting would depend on a modelling choice. $n = 307$.

#### B.8 The most conservative covariate model

Three specifications were fitted for the age effect on mean curvature, each adding a covariate that could carry part of it.

**Table B3:** Variance in mean curvature attributable to age under the total, direct and conservative covariate specifications.

| model | covariates | $R^2$ for age |
|---|---|---|
| total | sex | 0.279 |
| direct | sex, total streamline weight | 0.179 |
| conservative | sex, total weight, streamline count | 0.151 |

The conservative model is reported for completeness rather than as the preferred estimate. Total streamline weight and streamline count are both partly consequences of ageing rather than nuisance variables independent of it, so adjusting for both removes some of the effect under study along with the artefact. The total and direct estimates bracket the quantity of interest; the conservative estimate is a lower bound.

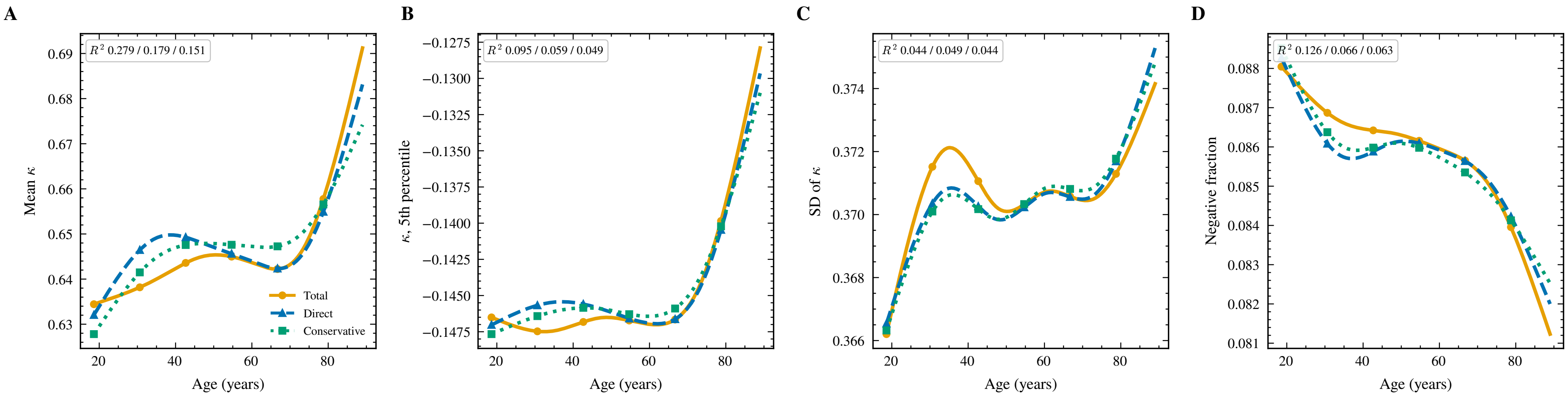

**Figure B6:** The shape of the age course is preserved across covariate specifications; only the variance attributed to age changes. (A) Mean curvature under the three specifications. (B) Fifth percentile of the edge curvature distribution. (C) Standard deviation of the edge curvature distribution. (D) Fraction of negatively curved edges. Total adjusts for sex; direct adds total streamline weight; conservative adds streamline count. The latter two are partly consequences of ageing rather than nuisance variables independent of it, so the conservative estimate is a lower bound. Statistics in each panel give the age $R^2$ under the three specifications in the order total, direct, conservative. $n = 307$.

##### B.9 The complete network block matrix

Seven of the 23 testable blocks survived correction and are reported in the main text. The remaining sixteen are given in Table B1 for completeness. Five blocks carried fewer than 20 retained edges and were excluded from modelling rather than modelled and corrected; excluding them before correction rather than after avoids spending the correction budget on estimates that could not have been reliable.

**Table B4:** All 23 modelled network blocks, ordered by age effect.

| Block | Kind | Edges | Beta | t | p (FDR) |
|---|---|---|---|---|---|
| Cont-Default | inter | 257 | $-4.20 \times 10^{-4}$ | -5.65 | $< 1 \times 10^{-4}$ |
| Vis-DorsAttn | inter | 68 | $-4.71 \times 10^{-4}$ | -3.77 | $9.0 \times 10^{-4}$ |
| Limbic-Cont | inter | 23 | $-7.06 \times 10^{-4}$ | -3.20 | 0.006 |
| Limbic-Default | inter | 60 | $-4.54 \times 10^{-4}$ | -2.50 | 0.042 |
| Default-Default | intra | 199 | $-2.09 \times 10^{-4}$ | -2.09 | 0.107 |
| Vis-Cont | inter | 57 | $-1.47 \times 10^{-4}$ | -1.84 | 0.171 |
| SalVentAttn-Default | inter | 195 | $-8.56 \times 10^{-5}$ | -1.45 | 0.339 |
| DorsAttn-Cont | inter | 124 | $-9.27 \times 10^{-5}$ | -1.12 | 0.464 |
| SomMot-Default | inter | 115 | $-4.97 \times 10^{-5}$ | -0.88 | 0.583 |
| SomMot-Cont | inter | 114 | $-4.58 \times 10^{-5}$ | -0.81 | 0.599 |
| SomMot-SomMot | intra | 195 | $-8.02 \times 10^{-5}$ | -0.64 | 0.665 |
| Cont-Cont | intra | 80 | $-7.28 \times 10^{-5}$ | -0.44 | 0.803 |
| Vis-Vis | intra | 204 | $-3.43 \times 10^{-5}$ | -0.28 | 0.855 |
| SalVentAttn-Cont | inter | 138 | $-1.51 \times 10^{-5}$ | -0.18 | 0.894 |
| DorsAttn-Default | inter | 159 | $+2.11 \times 10^{-6}$ | +0.03 | 0.973 |
| DorsAttn-DorsAttn | intra | 82 | $+5.24 \times 10^{-5}$ | +0.32 | 0.855 |
| Vis-Default | inter | 164 | $+4.50 \times 10^{-5}$ | +0.77 | 0.601 |
| SomMot-DorsAttn | inter | 199 | $+5.26 \times 10^{-5}$ | +0.92 | 0.583 |
| SalVentAttn-SalVentAttn | intra | 73 | $+2.04 \times 10^{-4}$ | +1.16 | 0.464 |
| Vis-Limbic | inter | 57 | $+1.10 \times 10^{-4}$ | +1.34 | 0.379 |
| SomMot-SalVentAttn | inter | 202 | $+2.70 \times 10^{-4}$ | +3.89 | $7.1 \times 10^{-4}$ |
| DorsAttn-SalVentAttn | inter | 122 | $+3.45 \times 10^{-4}$ | +5.09 | $< 1 \times 10^{-4}$ |
| Vis-SalVentAttn | inter | 29 | $+1.35 \times 10^{-3}$ | +7.64 | $< 1 \times 10^{-4}$ |

##### B.10 Analyses that returned nothing

Two further analyses are recorded here because a reader may wonder whether they were attempted.

**Polarisation of the existing curvature gradient.** If ageing amplified existing geometric differences between nodes, baseline curvature and the age coefficient would correlate positively; if it compressed them, negatively. Neither held ($\rho = -0.108$, $p = 0.129$). The dispersion of nodal curvature across nodes rose slightly, from 0.043 in the youngest tercile to 0.048 in the oldest, which is in the direction of polarisation but not distinguishable from noise.

**Conventional nodal properties of the decreasing set.** Degree, participation coefficient, clustering, betweenness and baseline curvature were compared between the decreasing and increasing sets, and none differed significantly (all $p > 0.11$). The point estimates were consistently in the same direction, with the decreasing nodes showing lower degree ($d = -0.39$), lower participation ($d = -0.56$) and lower clustering ($d = -0.38$), which would place them peripherally rather than as hubs. With 10 nodes against 57 this comparison could detect only effects above $d = 0.97$ at 80% power, so the null result carries no evidential weight in either direction and the point estimates should not be interpreted.

#### B.11 Cohort, prediction and nodal properties

Three figures supporting statements made in the manuscript without occupying space there.

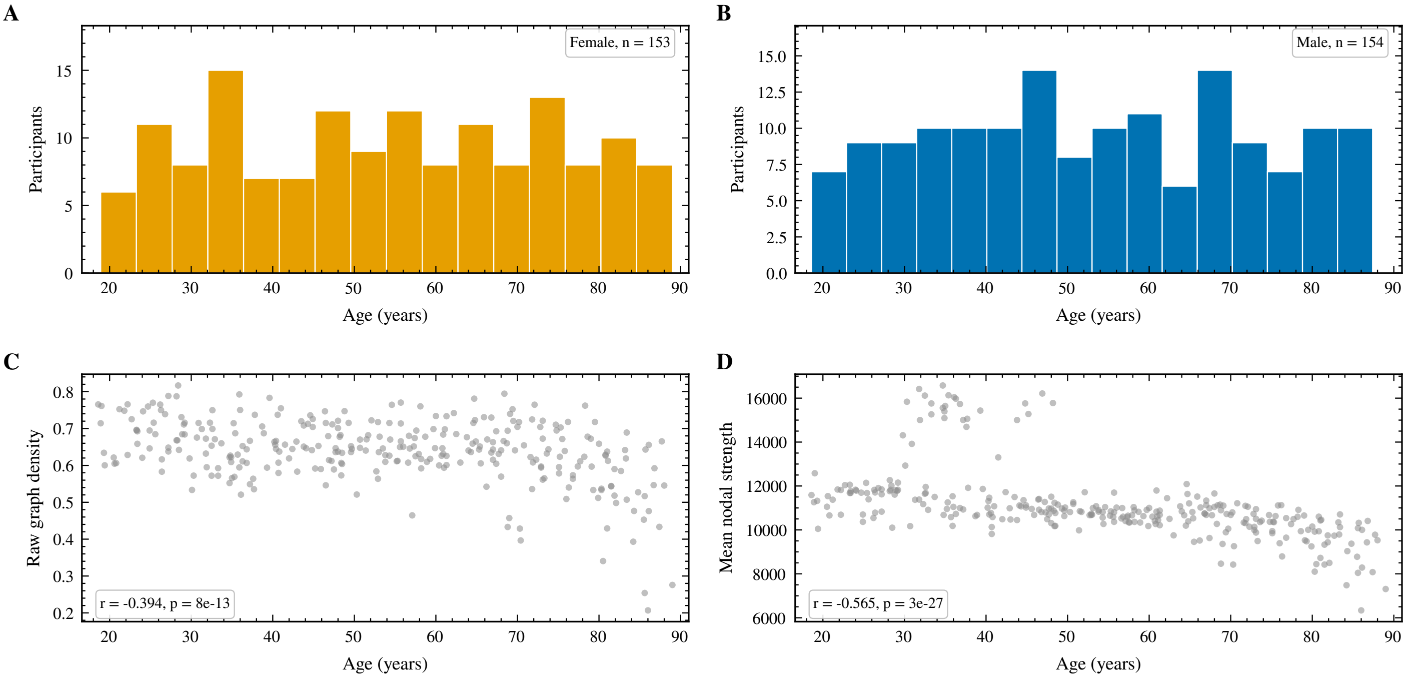

**Figure B7:** Cohort composition and the two quantities that motivate harmonisation and covariate adjustment. (A) Age distribution of female participants. (B) Age distribution of male participants. (C) Raw graph density against age, before harmonisation. (D) Mean nodal strength against age. Both raw density and total strength decline with age, which is why every graph is reduced to an identical edge count and why total strength is carried as a covariate.

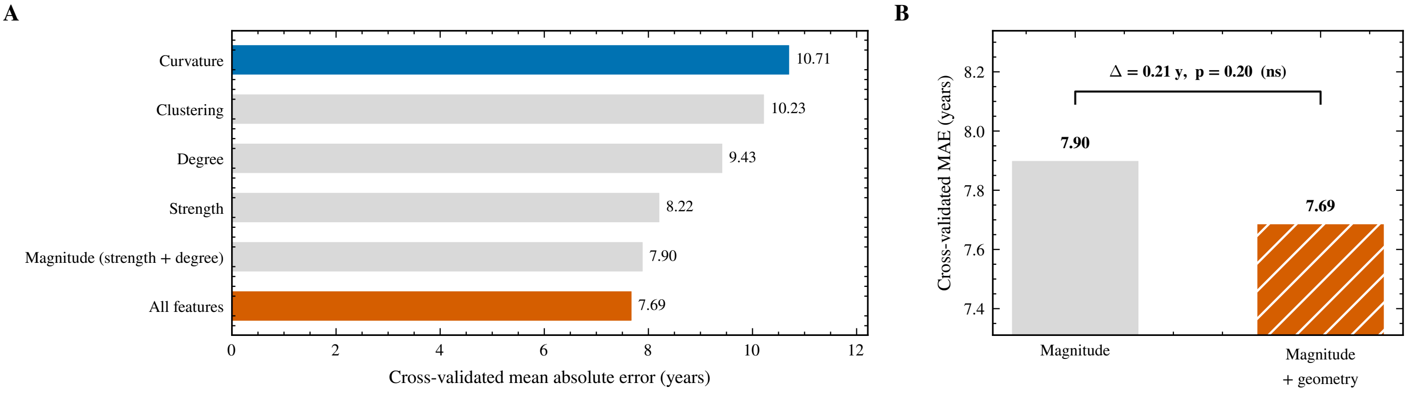

**Figure B8:** Curvature does not outperform magnitude at predicting age and adds no significant incremental accuracy. (A) Cross-validated mean absolute error of ridge regression predicting age from each feature block, over ten folds. (B) The two nested models compared directly on an expanded error scale, with the change in error and the p value of a sign-flip permutation test on the paired per-participant errors. This negative result is reported deliberately: the claim of this work is not predictive accuracy.

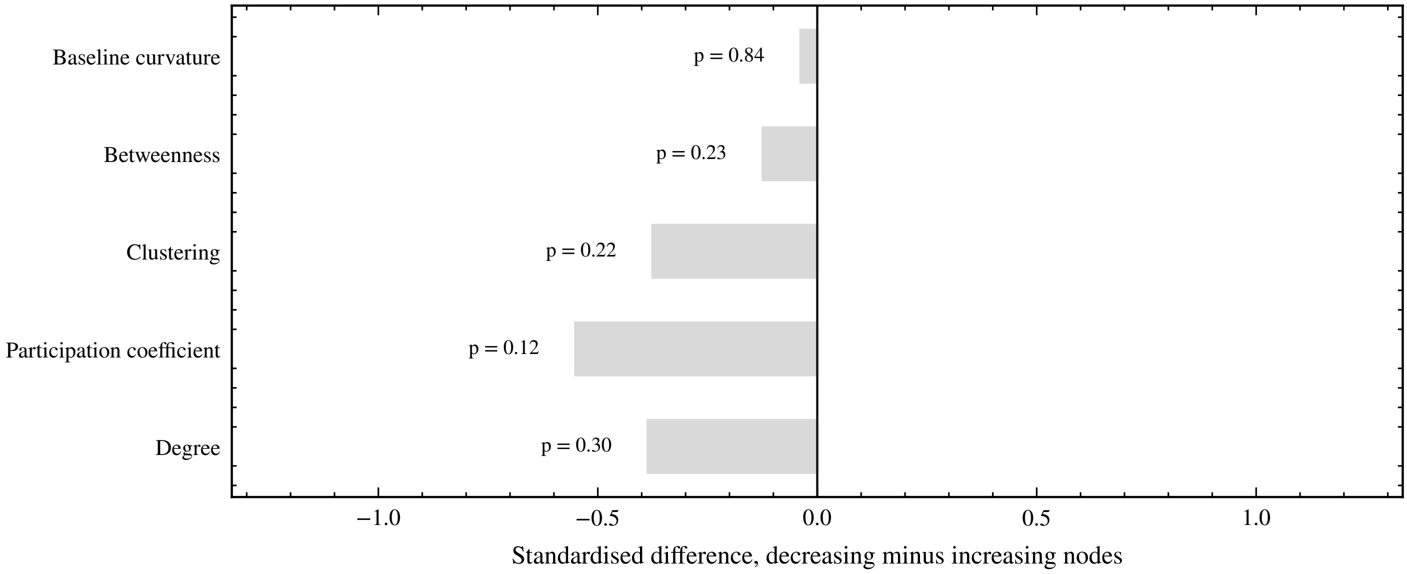

**Figure B9:** The decreasing nodes are not distinguishable from the increasing nodes by any classical nodal property. Degree, participation coefficient, clustering, betweenness and baseline curvature were compared between the two sets and no comparison reached significance. With 10 nodes against 57 the comparison could detect only effects above $d = 0.97$ at 80% power, so the point estimates should be read as descriptive.

### Annex C. Extended tractography and connectome methods

The Methods section of the manuscript describes the processing chain at the level a reader needs to follow the argument. This annex names every piece of software, every command-line option and the estimator behind each step, so that the reconstruction can be repeated exactly.

The Cam-CAN data are obtained from https://cam-can.mrc-cbu.cam.ac.uk/dataset/ free of charge, following the access procedure described there. Everything below applies to the released data as distributed, so a reader with access can reproduce the connectivity matrices analysed in the manuscript without further information from us.

#### C.1 Sample and acquisition

Diffusion data came from Stage 2 of the Cam-CAN CC700 release (Shafto et al., 2014; Taylor et al., 2017). Acquisition used a 3 T Siemens TIM Trio (32-channel head coil) and a twice-refocused spin-echo EPI sequence chosen by the original investigators for its intrinsic reduction of eddy-current distortion (Reese et al., 2003). The scheme comprises two non-zero shells, $b = 1000$ and $b = 2000\ \mathrm{s\,mm^{-2}}$, with 30 non-collinear directions each, plus three $b = 0$ volumes, for 63 volumes in total. Geometry was 66 axial slices, 2 mm isotropic voxels, FOV $192 \times 192$ mm, matrix $96 \times 96$, TR = 9100 ms, TE = 104 ms, partial Fourier 7/8, GRAPPA factor 2 with 36 reference lines, one average, acquisition time 10 min 2 s; these are collected in Annex C, Table C1. Phase encoding was anterior–posterior with no reverse phase-encoded acquisition. Shell structure, phase-encoding direction and total readout time were read per participant from the b-value table and the BIDS JSON sidecar rather than assumed.

The anatomical reference was the participant's own T1w MPRAGE from the same session. Processing ran on a single workstation, specified in Annex C, Table C2, under Python 3.11, with four participants processed concurrently and seven threads per participant.

#### C.2 Software

MRtrix3 3.0.8 (Tournier et al., 2019) provided the diffusion-specific steps and called FSL 6.0.7.22 (Jenkinson et al., 2012) and ANTs 2.6.5 (Avants et al., 2011) for distortion correction, registration, bias correction and tissue segmentation; bias correction used `N4BiasFieldCorrection` 2.6.5. Skull stripping used HD-BET 2.0.1 (Isensee et al., 2019) on GPU. Susceptibility correction used the Synb0-DisCo v3.1 container (Schilling et al., 2019, 2020) under Singularity, with `prepare_input.sh` patched to replace the missing `nu_correct` binary with ANTs `N4BiasFieldCorrection`. Surface reconstruction and the native-space Schaefer parcellation used FreeSurfer 8.2.0 (Fischl, 2012). Matrix assembly used NumPy 2.4.4 and nibabel 5.4.2.

#### C.3 Preprocessing

###### Conversion.

`mrconvert` embedded the FSL-format gradient table and the JSON metadata into a single `.mif` image, resolving the gradient-orientation convention once at the start.

###### Denoising.

`dwidenoise` performs Marchenko–Pastur PCA denoising over sliding $5 \times 5 \times 5$ patches (Veraart et al., 2016). For a patch matrix of $M$ voxels by $N$ volumes with ratio $\gamma = M/N$, the eigenvalues of pure noise are bounded by

$$\lambda_{\pm} = \sigma^2 \left(1 \pm \sqrt{\gamma}\right)^2,$$

so components with eigenvalue above $\lambda_+$ are retained as signal, the remainder are discarded, and $\sigma$ is estimated from the discarded bulk. This step precedes every interpolating operation, since the estimator assumes independent noise across voxels. The noise map and the residual image were kept for quality control.

###### Gibbs ringing.

`mrdegibbs -nshifts 20` re-evaluates each line of the image on a grid of subvoxel shifts $s/(2K)$ and keeps, per voxel, the shift that minimises local total variation,

$$D(s) = \sum_n \left|I_s(n+1) - I_s(n)\right|,$$

then interpolates back to the original grid (Kellner et al., 2016). The acquisition uses partial Fourier 7/8, which reduces the effectiveness of the correction but does not invalidate it.

###### Motion and eddy currents.

`dwifslpreproc -rpe_none -pe_dir AP` wrapped FSL `eddy` with the options `–slm=linear –repol –cnr_maps –residuals –data_is_shelled –dont_peas`. `eddy` models the diffusion signal as a Gaussian process over q-space, predicts each volume from all others, and estimates a rigid motion parameter set and an eddy-current field per volume by maximising agreement with that prediction (Andersson and Sotiropoulos, 2016). `–repol` replaces slices whose observed signal departs from the prediction by more than four standard deviations (Andersson et al., 2016). `–slm=linear` adds a linear second-level model appropriate for direction schemes that do not sample the sphere densely, and `–dont_peas` disables post-eddy alignment of shells along the phase-encoding direction, which fails on single-phase-encoding multi-shell data in the FSL version used here. Gradient directions were rotated with the estimated motion parameters and re-exported for all downstream steps.

###### Susceptibility distortion.

In EPI, off-resonance produces a displacement along the phase-encoding direction of

$$d(x) = f(x)\, T_{\text{readout}},$$

with $f$ the field offset in Hz and $T_{\text{readout}}$ the total readout time. With no reverse phase-encoded volume, $f$ cannot be estimated from the diffusion data alone. Synb0-DisCo predicts an undistorted $b = 0$ image from the T1w scan with a deep network trained on paired distorted and undistorted data (Schilling et al., 2019, 2020). The container was run with `–notopup`, and its output `b0_u` was merged with the measured mean $b = 0$ image computed from the eddy-corrected series. `topup` then estimated the field from that pair, with an `acqparams` file that gives the measured image its true readout time and the synthetic image a readout time of zero, that is, effectively infinite bandwidth. `applytopup –method=jac –inindex=1` applied the resulting displacement field to the eddy-corrected series with Jacobian intensity modulation,

$$I_{\text{corr}}(x) = I(x + d(x)) \left|1 + \frac{\partial d(x)}{\partial x}\right|.$$

Note the ordering: `eddy` runs before the field is known, and the susceptibility correction is applied afterwards as a separate resampling, rather than being passed to `eddy` through its `–topup` argument. Both corrections are therefore applied, but not jointly estimated. Two properties of this protocol limit what that separation can cost. The twice-refocused spin-echo sequence cancels eddy-current distortion at acquisition (Reese et al., 2003), so the volume-specific field that `eddy` estimates is dominated by head motion rather than by residual eddy currents, and the synthetic $b = 0$ image that constrains TOPUP is derived from the T1w scan rather than from the diffusion series, so its estimate does not depend on `eddy` having run first. The susceptibility field itself is static across volumes, which is what makes it admissible to apply it once, after motion correction, rather than per volume. The cost of the separation is therefore one additional interpolation of the diffusion series and not a volume-specific geometric error.

###### Bias field.

`dwibiascorrect ants` applied N4 (Tustison et al., 2010), which in the log domain treats the measured image $v$ as $v = u + b$ with $b$ a smooth B-spline field, and iterates between histogram sharpening and field fitting. The estimated field was retained for quality control.

###### Brain mask.

HD-BET was run on the reoriented T1w image; the resulting mask was transferred to diffusion space by a 6-degree-of-freedom FLIRT registration with a normalised mutual information cost, applied with nearest-neighbour interpolation, and binarised. All participants were masked by this route.

##### C.4 Fibre orientation distributions

Response functions for the three tissue classes were estimated with `dwi2response dhollander`, which selects representative single-fibre white matter, grey matter and CSF voxels without a tissue prior (Dhollander et al., 2016, 2019). Fibre orientation distributions were then estimated with `dwi2fod msmt_csd` (Jeurissen et al., 2014), which models the signal at gradient direction $\mathbf{u}$ and b-value $b$ as a sum of per-tissue convolutions,

$$S(b, \mathbf{u}) = \sum_t \sum_{l,m} f_{t,lm}\, R_{t,l}(b)\, Y_{lm}(\mathbf{u}),$$

with $R_{t,l}$ the rotational harmonic coefficients of the response of tissue $t$ and $Y_{lm}$ the spherical harmonic basis. Coefficients are obtained by non-negativity constrained least squares,

$$\min_{\mathbf{f}} \|\mathbf{Af} - \mathbf{S}\|^2 \quad \text{subject to} \quad \mathbf{Bf} \geq 0,$$

the constraint being applied to FOD amplitudes on a dense direction set. The diffusion scheme sampled three shells ($b = 0$, 1000 and 2000 $\mathrm{s\,mm^{-2}}$) with 3, 30 and 30 volumes respectively, and the maximum spherical harmonic order was set from those data: the white matter ODF was estimated at $l_{\max} = 8$, that is 45 spherical harmonic coefficients, read back from the image header rather than assumed, while the grey matter and CSF ODFs are isotropic ($l_{\max} = 0$) by construction. That order is supported by the two non-zero shells jointly and not by either alone, since 45 coefficients cannot be recovered from 30 directions at a single b-value, whereas the multi-tissue fit draws on all 63 volumes to solve for 47 unknowns across the three tissue compartments. `mtnormalise` then estimated a smooth multiplicative field in the log domain and rescaled the three tissue ODFs so that their summed volume fractions are spatially uniform (Raffelt et al., 2017), which is the condition under which SIFT2 weights and the $\mu$ coefficient are comparable across participants. As a quality-control measure, the masked mean of the $l = 0$ term of the white-matter FOD was recorded for every participant.

##### C.5 Anatomically constrained tractography

A five-tissue-type (5TT) image was generated from the T1w scan with `5ttgen fsl -premasked -nocrop`, which runs FSL FAST for cortical grey matter, white matter and CSF partial volumes (Zhang et al.,

2001) and FSL FIRST for subcortical grey matter (Patenaude et al., 2011), taking the HD-BET brain as input so that FIRST is not confronted with extracerebral tissue. All participants were segmented by this route, so the tissue partial volumes driving the anatomical constraints are of the same kind throughout the cohort.

The 5TT image was carried into diffusion space with the same transform used later for the parcellation. The transform was computed once, by registering the skull-stripped T1w image to the mean $b = 0$ image with FLIRT at 6 degrees of freedom and a normalised mutual information cost, and converting the result to a world-space transform with `transformconvert …flirt_import`. FSL matrices are expressed in stored-voxel coordinates and are valid only for the exact pair of image geometries they were computed for, whereas MRtrix transforms are in world coordinates; all subsequent resampling used the world-space form through `mrtransform -linear`. The source image was verified to be a skull-stripped brain, defined as a nonzero fraction of the field of view between 4 and 30 per cent, before any registration was accepted. The grey-matter/white-matter interface used for seeding was derived from the coregistered 5TT image with `5tt2gmwmi`.

Whole-brain tractography used `tckgen -algorithm iFOD2` (Tournier et al., 2010). iFOD2 draws candidate paths as short circular arcs and accepts them by rejection sampling with probability proportional to the product of FOD amplitudes at sample points along the arc, which makes it less prone than first-order methods to cutting corners at sharp bends. Parameters were:

- `-select`, 5 000 000: streamlines accepted per participant
- `-cutoff`, 0.06: FOD amplitude termination threshold
- `-step`, 0.625 mm: integration step (0.31 voxel)
- `-angle`, 45°: maximum curvature per step
- `-minlength`, 5 mm: minimum accepted length
- `-maxlength`, 250 mm: maximum accepted length
- `-act`, `5tt_coreg.mif`: anatomically constrained tractography
- `-backtrack`, on: retract and resample on invalid termination
- `-crop_at_gmwmi`, on: truncate endpoints at the interface
- `-seed_gmwmi`, `gmwmi.mif`: seeding from the grey/white interface

Anatomical constraints follow Smith et al. (2012): streamlines must terminate at the grey-matter/white-matter interface or in subcortical grey matter, are rejected if they end inside white matter or enter CSF, and with `-backtrack` a streamline that reaches an invalid termination retreats and resamples rather than being discarded.

Streamline weights were then estimated with `tcksift2` (Smith et al., 2015), which assigns each streamline $f$ a non-negative weight $w_f$ such that the weighted streamline density in each fixel $v$ matches the fibre density implied by the FOD,

$$\mu \sum_{f \in v} w_f \, \ell_{f,v} \;\approx\; \mathrm{FD}_v,$$

where $\ell_{f,v}$ is the length of streamline $f$ within fixel $v$ and $\mu$ is a single global proportionality coefficient per participant. The weights minimise a cost that sums a per-fixel discrepancy term over all fixels and a regularisation term penalising deviation of $\log w_f$ from zero. SIFT2 was run with the same ACT image used for tracking, and $\mu$ and the coefficient trace were saved. Unlike SIFT, SIFT2 discards no streamlines, so the full five million are available with weights.

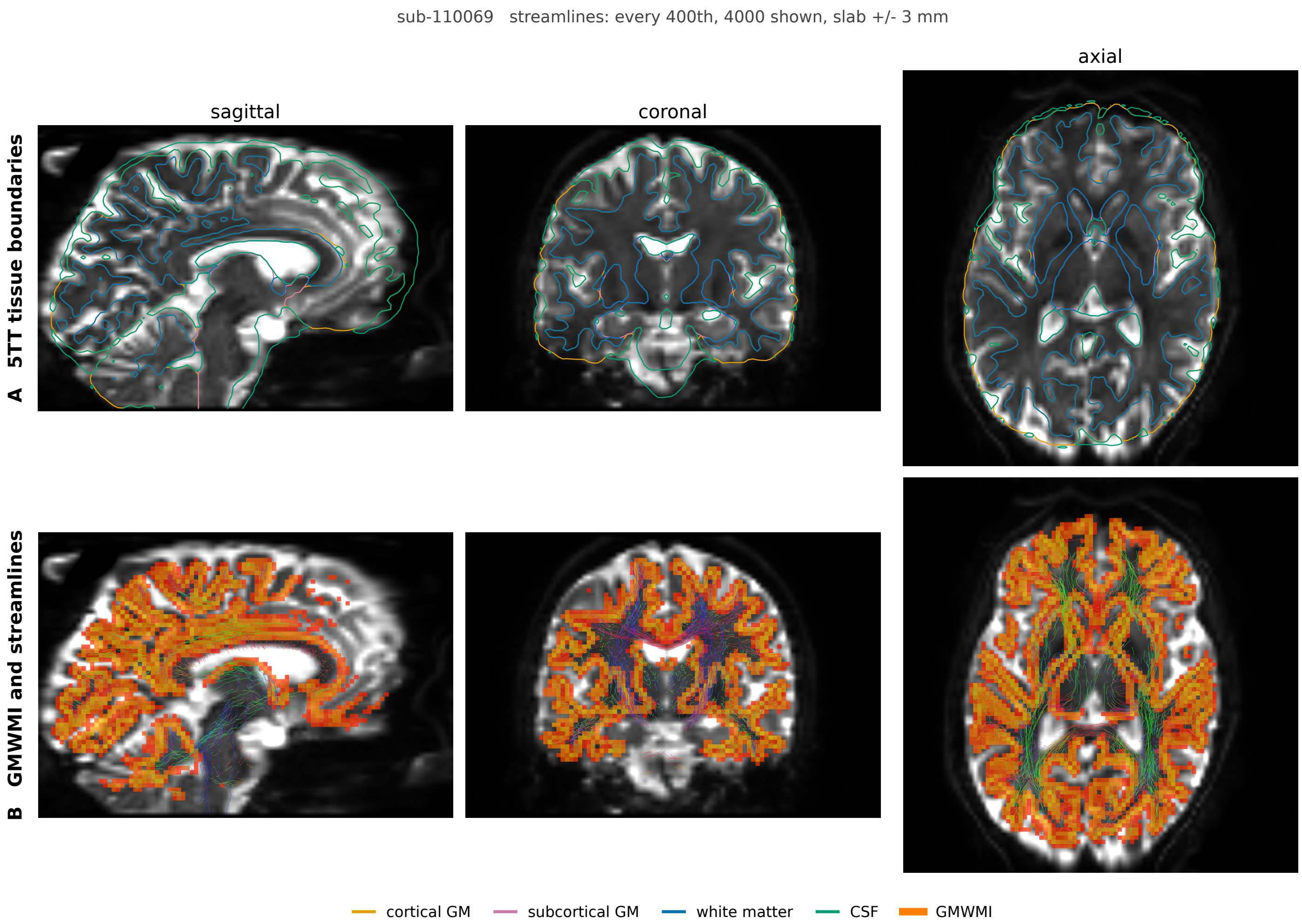

**Figure C1: Anatomically constrained tractography in one participant.** (A) Five-tissue-type segmentation, shown as tissue boundaries over the participant's diffusion-weighted image in sagittal, coronal and axial view. Cortical grey matter, subcortical grey matter, white matter and cerebrospinal fluid are outlined separately; these boundaries are what the tracking algorithm consults to accept or reject each streamline. (B) The grey-matter to white-matter interface, from which streamlines are seeded, with a sample of the resulting tractogram over the same three views. Streamlines are shown for a slab of a few millimetres and thinned by a large factor, so the display is a sample of the tractogram rather than the tractogram itself.

##### C.6 Parcellation

Nodes were the 200 cortical parcels of the Schaefer local-global parcellation in its 7-network variant (Schaefer et al., 2018; Yeo et al., 2011). Rather than warping the MNI-space volumetric atlas into each participant, the fsaverage annotation was projected onto each participant's own surface reconstruction and converted to a volumetric label image over the cortical ribbon with FreeSurfer (`mri_surf2surf` followed by `mri_aparc2aseg`), giving `Schaefer200_7Networks+aseg.nii.gz` in the participant's anatomical space. This route removes the template-to-subject affine and the template brain-versus-head mismatch from the parcellation path altogether.

FreeSurfer writes cortical codes as $1000 +$ colortable index on the left and $2000 +$ index on the right, with a per-hemisphere table for Schaefer, so connectome indices were obtained by

$$n(\mathbf{x}) = \begin{cases} a(\mathbf{x}) - 1000, & 1001 \le a(\mathbf{x}) \le 1100, \\ a(\mathbf{x}) - 2000 + 100, & 2001 \le a(\mathbf{x}) \le 2100, \\ 0, & \text{otherwise,} \end{cases}$$

giving left hemisphere parcels 1–100 and right hemisphere parcels 101–200 in the same order as the label file. The label volume was then resampled onto the mean $b = 0$ grid with `mrtransform -linear t1w_to_dwi_mrtrix.txt -template mean_b0.mif -interp nearest -datatype int16`, that is, for each diffusion-space voxel $\mathbf{y}$,

$$L_{\mathrm{DWI}}(\mathbf{y}) = L_{\mathrm{nat}}\left(\mathrm{NN}\left(\mathbf{A}_{\mathrm{nat}}^{-1}\mathbf{T}^{-1}\mathbf{A}_{b_0}\mathbf{y}\right)\right),$$

with $\mathbf{A}$ the voxel-to-world affines, $\mathbf{T}$ the T1w-to-diffusion rigid transform and NN nearest-neighbour rounding. The whole path involves a single interpolation of the label image, performed in world coordinates. Parcel completeness was recorded per participant as the set of expected indices absent from the resampled volume, and label identity was verified independently of completeness, since a permutation of parcel indices fills every node and would pass a completeness check unnoticed. The verification compared the fsaverage annotation colortable order against the label file from which node names were taken, confirmed that the observed cortical codes occupy the per-hemisphere ranges 1001–1100 and 2001–2100 rather than the shared-table range that would silently discard the right hemisphere, recomputed the label volume from its own source and compared it voxelwise with the volume on disk, and checked the sign of each parcel centroid's world $x$ coordinate against its nominal hemisphere. No participant failed any check.

##### C.7 Connectome construction

`tck2connectome` assigned each streamline endpoint to a parcel using a radial search of up to 4 mm from the endpoint, taking the closest parcel encountered, and wrote the per-streamline assignment pairs alongside the streamline-count matrix, with `-symmetric` and `-zero_diagonal`. Streamlines with an unassigned endpoint, that is, those whose radial search reached no parcel, contribute to no edge, and neither do streamlines whose two endpoints fall in the same parcel, which the zero diagonal removes. Both were quantified rather than left implicit. Across a random subsample of 79 participants, both endpoints reached a parcel for $0.741 \pm 0.039$ of streamlines; of the whole tractogram, $0.241 \pm 0.022$ connected a parcel to itself and $0.500 \pm 0.044$ contributed to an inter-parcel edge. The remaining $0.259 \pm 0.039$ had at least one endpoint outside any parcel, which is expected for a cortical parcellation: anatomically constrained tractography permits termination in subcortical grey matter, and the Schaefer atlas defines no node there. The inter-parcel fraction was computed for every participant from the count matrix, which gives the same quantity because the accumulation zeroes the diagonal, and the two routes agreed exactly.

Writing $(a_s, b_s)$ for the parcel pair assigned to streamline $s$, $w_s$ for its SIFT2 weight, $\ell_s$ for its length and $m_s^{(k)}$ for the along-streamline mean of scalar map $k$, and $\delta_s^{ij} = \mathbf{1}[\{a_s, b_s\} = \{i, j\}]$, the matrices are

$$\begin{aligned} C_{ij} &= \sum_s \delta_s^{ij}, & W_{ij} &= \sum_s w_s\, \delta_s^{ij}, \\ L_{ij} &= \frac{1}{C_{ij}} \sum_s \ell_s\, \delta_s^{ij}, & L_{ij}^{\mathrm{inv}} &= \frac{1}{C_{ij}} \sum_s \ell_s^{-1}\, \delta_s^{ij}, \\ M_{ij}^{(k)} &= \frac{1}{C_{ij}} \sum_s m_s^{(k)}\, \delta_s^{ij}, \end{aligned}$$

for $i, j \in \{1, \dots, 200\}$, with $W_{ij} = W_{ji}$ and $W_{ii} = 0$ by construction, and edges with $C_{ij} = 0$ set to zero rather than left undefined. The SIFT2-weighted matrix $W$ is the primary connectome used in all analyses. Scalar values $m_s^{(k)}$ were obtained with `tcksample -stat_tck mean` from the diffusion tensor maps (`dwi2tensor` with nonlinear least squares, then `tensor2metric` for FA, MD, AD and RD) and from any additional voxelwise maps available for the participant; streamline lengths came from `tckstats -dump`. Endpoint assignment was performed once per participant per atlas, and the remaining matrices were accumulated from the stored assignment pairs, which reproduces `tck2connectome` semantics without re-reading the tractogram for each weighting.

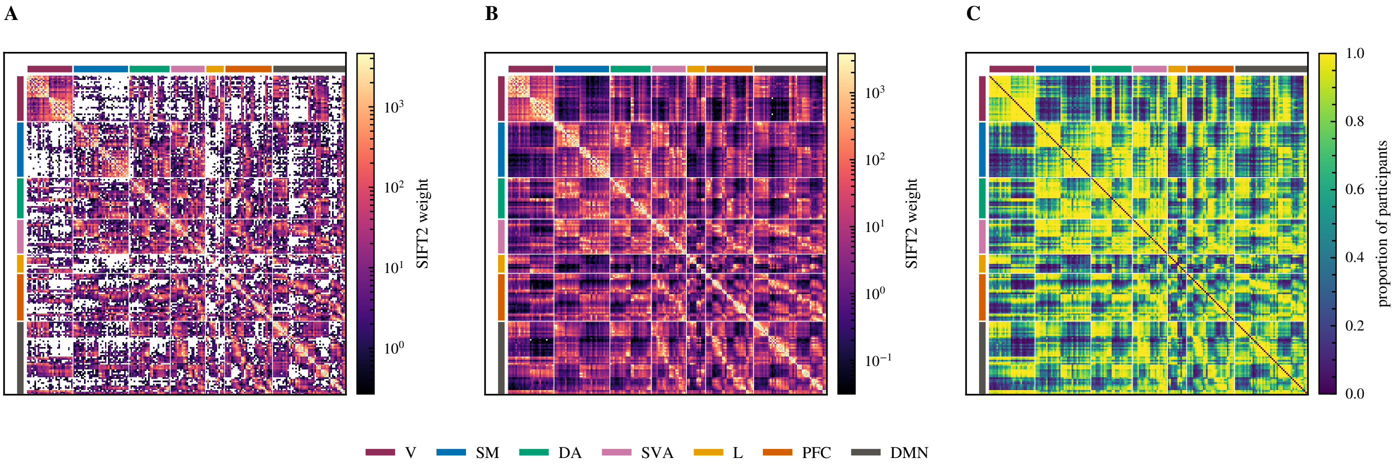

**Figure C2: Connectivity matrices at three levels.** Nodes are ordered by network and the coloured bars along the margins mark the seven canonical networks. (A) One participant chosen at random, on a logarithmic scale; white cells are node pairs with no reconstructed connection in that participant. (B) The mean across the cohort on the same scale, where the block structure of the parcellation becomes visible. (C) Edge consistency, the proportion of participants in whom each node pair carries a connection. This is the quantity the consistency mask thresholds: pairs connected in fewer than 60% of participants are excluded before harmonisation.

##### C.8 Quality control and exclusions

The following were recorded per participant and inspected before analysis: the noise map and denoising residuals; the estimated bias field; the masked mean of the $l = 0$ FOD term; the SIFT2 proportionality coefficient $\mu$; the nonzero fraction of the registration source image, used to confirm skull stripping; the number of empty parcels in the resampled parcellation; and the fraction of streamlines contributing to an inter-parcel edge. The SIFT2 coefficient was $2.47 \times 10^{-4} \pm 4.78 \times 10^{-5}$ (median $2.36 \times 10^{-4}$, range 1.61 to $4.49 \times 10^{-4}$), a coefficient of variation of 0.19 with a 2.8-fold spread between the extreme participants. Its absolute value carries no interpretation on its own, since it scales with the FOD amplitude and with the streamline count; what the narrow distribution shows is that multi-tissue intensity normalisation left the cohort on a common scale, which is the condition under which SIFT2 weights are comparable between participants. Every retained participant had all 200 parcels present in the resampled parcellation. Of the 312 datasets with complete diffusion input, five were excluded: three whose assignment fraction lay more than three standard deviations from the cohort mean (two below, one above), and two for which `tck2connectome` failed, leaving $n = 307$.

##### C.9 Provenance of the reported values

No placeholders remain. The quantitative figures in the text were measured rather than estimated, by two scripts kept with the analysis code:

- Assignment fractions, empty parcels: `measure_connectome_qc.py`
- SIFT2 $\mu$: `sift2_mu_stats.py`
- $l_{\max}$, shell structure: `mrinfo` on the FOD and preprocessed DWI
- Software versions: recorded on the processing host

Both scripts restrict their statistics to the analysis sample rather than to whatever is present on disk, and the assignment fraction is computed against the streamline count read from each tractogram header rather than against the nominal five million.

##### C.10 Tables

**Table C1:** Diffusion MRI acquisition parameters, Cam-CAN CC700 Stage 2 (Shafto et al., 2014; Taylor et al., 2017).

| **Parameter** | **Value** |
|---|---|
| Scanner | 3 T Siemens TIM Trio |
| Head coil | 32-channel |
| Sequence | twice-refocused spin-echo EPI (Reese et al., 2003) |
| Diffusion weighting | $b = 1000$ and $b = 2000\ \mathrm{s\,mm^{-2}}$ |
| Directions per shell | 30 non-collinear |
| Non-diffusion-weighted volumes | 3 |
| Volumes acquired | 63 |
| Voxel size | $2 \times 2 \times 2$ mm |
| Slices | 66, axial |
| Field of view | $192 \times 192$ mm |
| Acquisition matrix | $96 \times 96$ |
| Repetition time | 9100 ms |
| Echo time | 104 ms |
| Partial Fourier | 7/8 |
| Parallel imaging | GRAPPA factor 2, 36 reference lines |
| Averages | 1 |
| Acquisition time | 10 min 2 s |
| Phase-encoding direction | anterior–posterior |
| Reverse phase-encoded data | none acquired |
| Anatomical reference | T1w MPRAGE, same session |

Shell structure, phase-encoding direction and total readout time were read per participant from the b-value table and the BIDS JSON sidecar rather than taken from this table.

**Table C2:** Processing environment.

| **Component** | **Specification** |
|---|---|
| CPU | AMD Ryzen 9 5950X, 16 cores / 32 threads |
| Memory | 128 GB DDR4-3200 ($4 \times 32$ GB) |
| GPU | NVIDIA GeForce RTX 3090, 24 GB GDDR6X |
| Motherboard | ASUS ROG Crosshair VIII Dark Hero, AMD X570S |
| Working storage | Samsung 990 Pro 2 TB NVMe PCIe 4.0 |
| Archival storage | Seagate 20 TB SATA HDD |
| Operating system | Ubuntu 24.04.4 LTS |
| Runtime | Python 3.11 |
| Parallelism | 4 participants concurrently, 7 threads each |

GPU-bound steps (Synb0-DisCo, FSL ‘eddy‘, HD-BET) were serialised across workers by an inter-process semaphore, with at most three concurrent GPU jobs, so that concurrent participants could not exhaust the 24 GB of video memory.
